# Microbial Weathering Analysis of Anshun Tunbao Artifacts

**DOI:** 10.1101/2024.08.26.609815

**Authors:** Paierzhati Abudureyimu, Xiaoyan Luo, Chu Gui, Manchun Liu, Xining Su, Dingsong Lan, Zhi Chen, Jerome Rumdon Lon, Jianfei Luo

## Abstract

Cultural heritages are the crystallization of human technology, culture and embody the efforts of many craftsmen in ancient times. Wooden cultural heritages are affected by their own materials, and are very susceptible to microbial invasion under suitable temperature and humidity conditions. This project mainly studies the microbial weathering of the core wood carving masks and wooden ancient stage in Anshun Tunpu cultural heritage, and uses scanning electron microscopy, high-throughput sequencing and traditional culture methods to analyze the characteristics of their microbial communities, and finds that the bacteria represented by Pseudomonas, Paenibacillus and Stenotrophomonas, and the fungi represented by Cladosporium, Alternaria and Aspergillus, are the core microorganisms shared by indoor and outdoor cultural heritage. The dominant genera such as Pseudomonas, Stenotrophomonas, and Cladosporium showed lignocellulose deradation ability. By cultivating insect eggs extracted from sampled specimens and analyzing the correlation with surface microbiota, we verified the role of insect eggs as vectors in dispersing key microbial communities. We speculate that these insects are likely to be Anobiidae family. Furthermore, by comparing the microbial compositions under different climatic conditions, we validated the influence of temperature, humidity, vegetation diversity, and microbial intrinsic structures on microbial growth. Therefore, monitoring the surrounding environment is crucial in microbial weathering studies and cultural heritage preservation efforts. This study pioneered the investigation of microbial weathering aspects on unique heritage of the Tuenbao culture, emphasizing the importance of preserving distinctive ethnic cultures. Additionally, it presented a case study on the microbial weathering of wooden artifacts in karst landscape environments.

## 1. Introduction

The Tunbao Culture, a distinctive regional culture, embodies the rich history and ethnic memories of Anshun, Guizhou. Its origins can be traced back to the early Ming Dynasty military garrison system. In the 14th year of the Hongwu reign (1381) during the Ming Dynasty, Emperor Zhu Yuanzhang implemented a series of military and administrative measures in order to consolidate control over the southwestern regions, including the establishment of military outposts and the implementation of the garrison system in areas such as Guizhou.The implementation of the system led the majority of the garrison soldiers and their families, who hailed mainly from Jiangnan, to carry their hometown’s cultural traditions and production techniques across vast distances to the remote mountains of Guizhou. Here, they not only preserved the lifestyle habits and cultural features of Jiangnan but also adapted and innovated based on Guizhou’s natural environment and social conditions, giving rise to a unique regional culture in architecture, clothing, language, folk arts, economic models, religious beliefs, and more - known as the Tunbao Culture. Therefore, the Tunbao Culture is lauded as a "living fossil" of the Ming Dynasty legacy.

As an integral part of the Tunbao Culture, local opera plays a crucial role in cultural inheritance and serves as a significant component of the spiritual life of the Tunbao people. The masks used in local opera, also known as "lian zi," constitute an indispensable element in the performances. Anshun local opera originated from the regional opera genre "Yiyang Qiang" in the Yiyang area of Jiangxi during the late Yuan and early Ming dynasties. The masks sampled in this study include "lian zi" originating from the Yuan Dynasty.These masks, all crafted from wood, exhibit exquisite carving techniques and rich color expressions, showcasing the Tunbao people’s reverence for heroic figures and their steadfast dedication to traditional culture.

The Yunshan Tunxilou Theatre, built during the Qing Dynasty, is part of the Yunshan Tun Ancient Architectural Complex in Anshun City, Guizhou Province. Initially established as a military garrison, it was approved by the State Council in 2001 as one of the fifth major historical and cultural sites protected at the national level. Serving as a venue for local opera performances, the Yunshan Tunxilou Theatre has witnessed numerous theatrical productions and stands as an important medium for the Tunbao Culture. The architectural style of the theatre blends military defense with folk beliefs, reflecting the wisdom and resilience of the Tunbao people.

Biodegradation generally refers to the various life activities of organisms indirectly to the surrounding environment, and biodegradation is irreversible. Microorganisms, including bacteria, fungi, algae, lichens and other living organisms, these microbial life activities are harmful to wooden cultural heritage. The microbial corrosion of cultural heritage is a spontaneous, long-term process, microbes in process of enzymes, organic acids and other secretions will erode the surface of wooden cultural heritage to make it discolouration, degradation of wooden cultural heritage of the main components and thus reduce the strength of the cultural heritage.

The cell wall of wood is mainly composed of organic materials such as cellulose, hemicellulose, and lignin. Due to the composition of wood itself, it is highly susceptible to erosion and degradation by fungi, bacteria. Based on the different aspects of wood degradation by fungi, they can be categorized into three types: decay fungi that degrade components of wood such as cellulose and lignin by secreting decomposing enzymes, discoloration fungi that change the color of wooden artifacts by staining the wood surface, and mold fungi that produce mycotoxins leading to mold formation in wood[1–3]. In different environments, various fungi exhibit different abundances through mutual antagonism or synergy, thereby deeply corroding wooden artifacts from various aspects. Bacterial damage to wooden artifacts is less common, with predominantly bacilli, filamentous bacteria, and micrococci possessing cellulase enzyme systems with high activity, capable of degrading non-crystalline cellulose. Moreover, bacteria can utilize their relatively small surface-to-volume ratio for assimilation or heterotrophy to degrade lignin.

Unlike the mechanism of fungal corrosion of wooden artifacts, bacteria erode wood by attaching sugar externally to the surface of wood cells. Bacteria in saturated wood can grow and reproduce using the main components of the wood cell wall as a carbon source[4], causing damage to progress to the middle lamella and eventually leading to secondary wall decomposition.

Based on their mechanisms of action and the morphology of wood diseases, wood-rot fungi can be primarily classified into white rot fungi belonging to Basidiomycota, brown rot fungi, and soft rot fungi belonging to Ascomycota[5]. White rot fungi degrade lignin to CO2 through oxidation-reduction processes, leaving cellulose behind, resulting in white rot. Brown rot fungi secrete carbohydrate-active enzymes (CAZymes) to hydrolyze cellulose or hemicellulose in wood into glucose[6], leaving black lignin, causing brown rot. Soft rot fungi first invade the surface of wood through hyphae, metabolizing carbohydrates to soften the wood and gradually degrade it, while also secreting enzymes to act on the secondary wall. Additionally, soft rot fungi exhibit adaptability to extreme environments[7]. Discoloration fungi cause wood discoloration by invading the wood surface with hyphae, indirectly providing favorable conditions for other microorganisms to degrade wood[8]. Mold growth and metabolism not only secrete enzymes to degrade wood components but also produce acidic secretions to acidify the wood surface, accelerating degradation by other microorganisms. Among molds, species such as Penicillium, Aspergillus, and Fusarium pose the most serious microbial damage to wood[9]. Based on the corrosion mechanisms of bacteria on wooden artifacts, they can be categorized into eroding bacteria capable of forming erosion grooves on the inner cavity surface of wood under anaerobic conditions, tunneling bacteria that degrade lignin under aerobic conditions to soften wooden artifacts[10], and hollowing bacteria that adhere to the inner cavity surface through extracellular mucilage, leading to cavitation of wood cell walls[11].Currently, a variety of biotechnologies such as traditional culture methods, denaturing gradient gel electrophoresis (DGGE), scanning electronic microscopy (SEM), and high-throughput sequencing can be used to identify microorganisms on the surface of artifacts. Sabatini et al[12]. employed traditional culture methods and microscopic observation to screen and evaluate the fungi present on 64 cultural heritage artifacts collected from the Montefeltro region and adjacent areas to understand the microbial involvement in relevant biological deterioration. Arenz et al[13]. conducted an analysis of fungal diversity in wood and soil environments in the Antarctic region by combining traditional culture methods with DGGE technology, identifying four predominant fungal groups dominated by filamentous ascomycetes. Han et al[14]. conducted an analysis of fungal diversity in wood and soil environments in the Antarctic region by combining traditional culture methods with DGGE technology, identifying four predominant fungal groups dominated by filamentous ascomycetes. Han et al. analyzed the fungal communities of the Nanhai No. 1 shipwreck and its environment, utilizing SEM, lightmicroscope(LM), and high-throughput sequencing techniques to identify the dominant phylum of Ascomycota and ten predominant genera of Fusarium. Beata et al[15]. combined field emission scanning electron microscopy(FESEM), metabolomics analysis, and high-throughput sequencing to observe the degree of degradation of artifacts, analyze organic acids and melanin secreted during microbial growth and reproduction processes, and assess microbial diversity. Different research methods have different emphases to meet the diverse needs of research projects. For example, traditional cultivation methods are simple and effective in observing the physiological characteristics of microorganisms, SEM provides a more intuitive observation of microbial presence on sample surfaces and their degradation levels, high-throughput sequencing allows for a more comprehensive understanding of microbial community characteristics. These methods are widely applied due to their inherent advantages.

Different microorganisms perform distinct functions; for instance, filamentous microorganisms can utilize hyphae growth to extend into the interior structure of wood, providing attachment points for other microorganisms. On the other hand, non-filamentous microorganisms (such as Actinobacteria and Proteobacteria) can form biofilms that facilitate the aggregation of diverse microorganisms, thereby maintaining moisture and nutrients[16]. The interactions among these microorganisms increase microbial diversity and deepen the extent of damage to cultural artifacts. Therefore, identifying the core microbial communities on the surfaces of wooden masks and wooden ancient stages and analyzing the competitive advantages of dominant bacterial genera are crucial for the preservation and protection of cultural artifacts. Based on high-throughput sequencing, we analyzed the microbial communities on the surfaces of various samples and classified the samples into indoor and outdoor categories based on the fluctuation sizes of various environmental factors.

We then conducted specific analyses on the microbial community characteristics for both indoor and outdoor artifacts in karst landform environments. The core microbial communities shared by indoor and outdoor artifacts under these conditions include bacteria represented by Pseudomonas, Paenibacillus, and Stenotrophomonas, as well as fungi such as Cladosporium, Alternaria, and Aspergillus. By investigating the physiological characteristics of dominant microbial genera, we inferred their competitive advantages. We conducted degradation experiments on isolated microorganisms to verify the degradation capabilities of strains including Pseudomonas, Stenotrophomonas, Fusarium, Trichoderma, Penicillium, and Cladosporium, exploring their mechanisms of damage to artifacts and the extent of such damage.

Furthermore, we analyzed the relationship between the eggs of Anobiidae family found during the isolation and purification process and microorganisms, revealing that insect eggs can serve as a medium for spreading microorganisms and expanding their colonization range. Finally, we compared the dominant microorganisms under different climatic conditions and it was observed that the growth of microorganisms in varied climates is closely linked to bacterial intrinsic structures, preferences for temperature and humidity, as well as the richness of surrounding vegetation. This study confirmed the variability of surface microorganisms on artifacts under different climates.

## 2. Materials and Methods

### 2.1 Sample collection

The team selected Anshun City, Guizhou Province, China, which possesses typical karst climate and topography, for sampling. Samples were collected from indoor heritage dominated by masks and outdoor heritage dominated by stages to investigate the commonalities and differences in microbial weathering of heritage under karst climate conditions. Five masks with microbial weathering characteristics (M1, M2, M3, M4, M5) were sampled, with three points selected on each mask: M1 (M1-1, M1-2, M1-3)(Figure 1A), M2 (M2-1, M2-2, M2-3)(Figure 1B), M3 (M3-1, M3-2, M3-3)(Figure 1C), M4 (M4-1, M4-2, M4-3)(Figure 1D), M5 (M5-1, M5-2, M5-3) (Figure 1E). A ancient theatrical stage with typical weathering features was sampled, with three points selected on the front (Q1, Q2, Q3)(Figure 2D), lateral (C1, C2, C3)(Figure 2E), and top (D1, D2, D3)(Figure 2A-C) of the stage for sampling. With the consent of the staff members who have resided on the stage for a long time, samples were collected from their facial skin and labeled as F, which would be a reference to explore the connection between microbial communities and human life activities.

**Figure 1.**
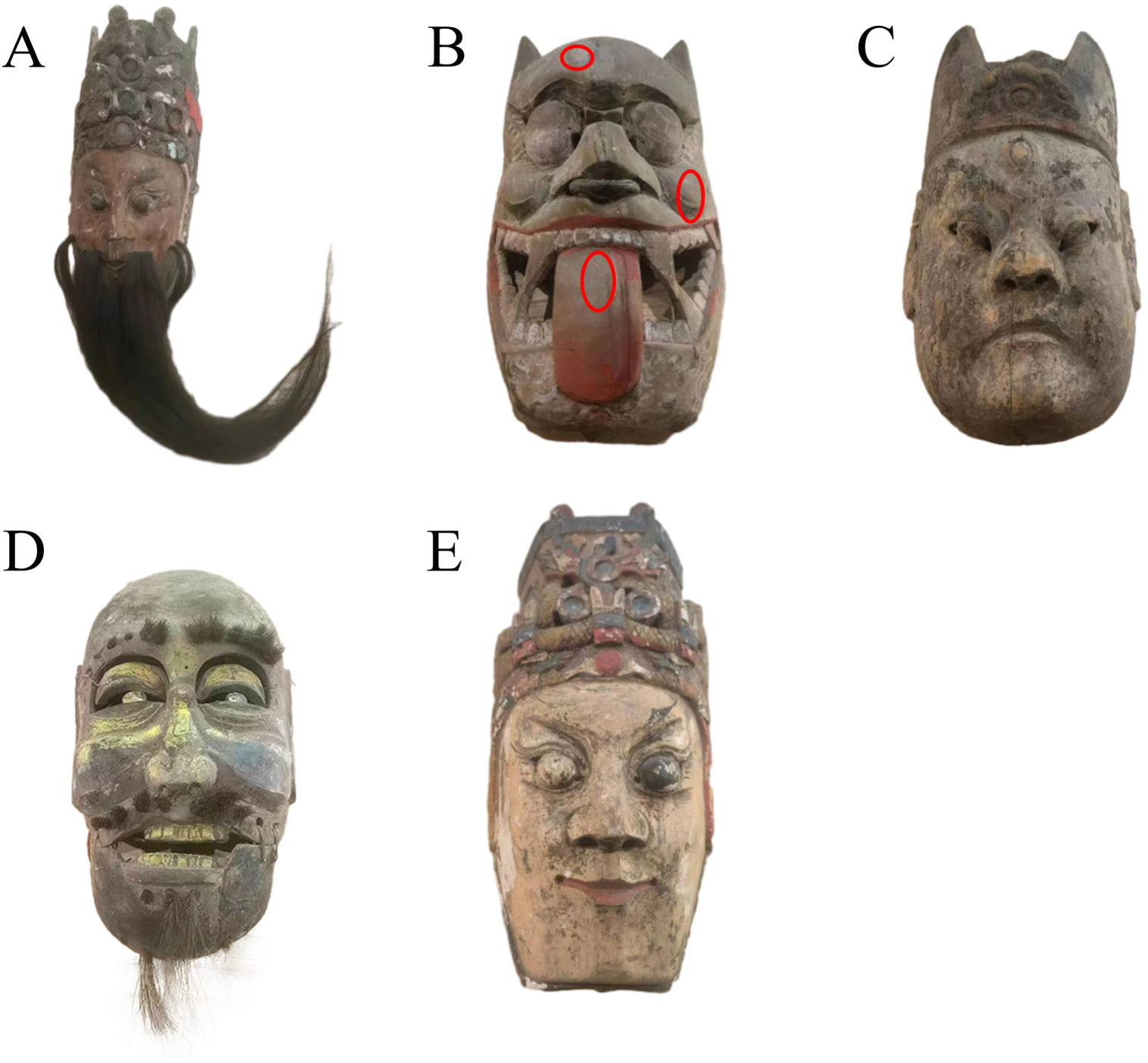
Characteristics of the mask samples. Sampling points on the masks. (A) Samples M1-1, M1-2, M1-3 collected from mask (M1); (B) Samples M2-1, M2-2, M2-3 collected from mask (M2); (C) Samples M3-1, M3-2, M3-3 collected from mask (M3); (D) Samples M4-1, M4-2, M4-3 collected from mask (M4); (E) Samples M5-1, M5-2, M5-3 collected from mask (M5); (F) Mask display wall. The material of hair on masks (A and D) is horsehair. The sampled masks are routinely hung on the display wall for preservation.

**Figure 2.**
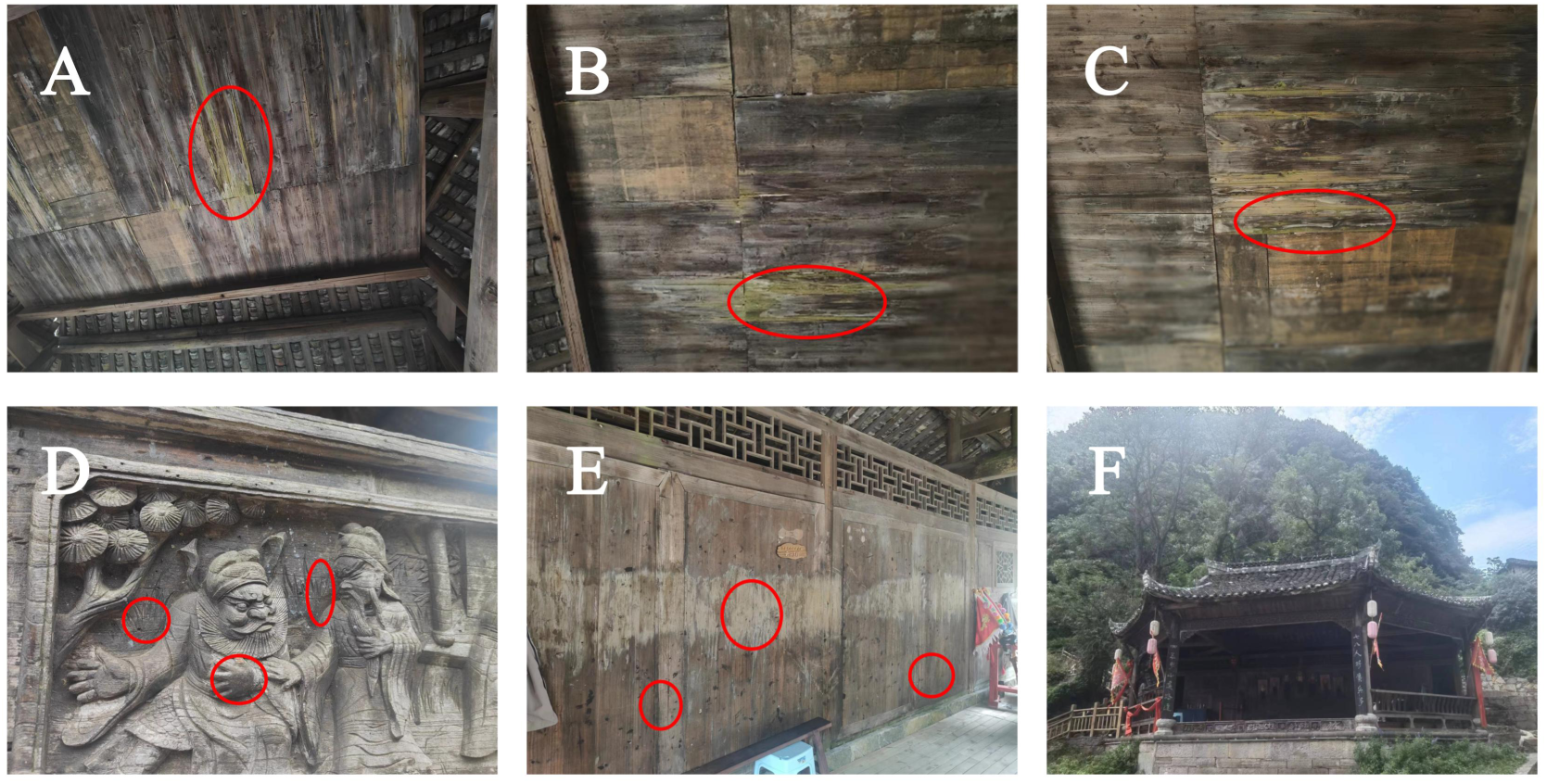
Characteristics of the theatre samples. Sampling points on the stage. (A, B, and C) Samples D1, D2, D3 collected from the top of the stage (D);(D) Samples C1, C2, C3 collected from the side of the stage (C);(E) Samples Q1, Q2, Q3 collected from the front side of the stage (Q);(F) Panoramic view of the stage. The marked areas in (A, B, C, D, and E) exhibit white rot and brown rot phenomena on wood after decay. The stage has good air circulation, and weeds are stacked in the front.

#### 2.1.1 Environmental factors recording

Using the SMART SENSOR AS817 thermometer and hygrometer (Dongguan Wanchong Electronic Products Co., Ltd.), relative humidity and temperature at the sampling sites were recorded. The environmental parameters for the masks remained relatively constant, with a temperature of 23.5°C and a relative humidity of 72.1±0.1. The stage environment exhibited minor fluctuations, with a temperature of 29.25±0.15°C and a relative humidity of 65.02±0.6. The moisture content of the wooden samples from the ancient stage was measured as 0 using the SMART SENSOR AS981 hygrometer. The moisture content of the wooden material of the masks was not measured to protect them. Environmental factors at the sampling sites were documented to facilitate resampling if necessary.

#### 2.1.2 Surface sampling protocol

Surface sampling will be conducted using sterile cotton swabs, which would be immersed in sterile M9 salt solution and rubbed on a 5 square centimeter area on the surface. Fresh M9 salt would be utilized for each sample to prevent cross-contamination between sampling points. Throughout all sampling stages, nitrile gloves would be worn to prevent contamination of samples with skin microbiota. Samples would be stored at 4°C for further cultivation or sequencing.

For sampling using carbon tape, a suitable length of carbon tape would be cut and gently placed on the surface of the artifact. The completed sampling carbon tape would be folded in half and placed in a sealed bag to prevent contamination of the tape surface. Samples would be stored at 4°C for subsequent observation of surface microbiota on artifacts using scanning electron microscopy.

### 2.2 Scanning Electron Microscope(SEM)

The sampled carbon tape will be affixed to a specimen stub and sputter-coated with gold for 30 seconds to enhance conductivity. Subsequently, the sample will be observed using a scanning electron microscope to identify areas exhibiting colony morphology characteristics for photographic documentation.

### 2.3 Isolation, purification and identification of microorganisms

The M9 salt sample solution will be vortexed at 150 rpm for 6 hours at 27°C to separate microorganisms from the swab. The resulting suspension will then be streaked onto non-selective culture media—LB agar for bacterial growth and YPD agar for fungal growth—and incubated at 27°C for 2-7 days to accommodate various growth states. Different strains will be isolated based on colony color and morphology, with repeated streaking on plates until pure cultures are obtained. For isolated and purified strains, the CTAB[17] extraction method will be employed for total microbial genomic DNA extraction to facilitate subsequent strain identification procedures.

#### 2.3.1 Bacterial Identification

For bacterial identification, the 16S rRNA V3-V4 region will be amplified using 2× Taq Master Mix (Sangon Biotech, B639295, China). Two universal bacterial 16S rRNA gene amplification PCR primers will be utilized: the forward primer 27F and the reverse primer 1492R. The reaction conditions are as follows: microbial DNA (10 ng/μl) 2 μl; forward primer for V3-V4 amplification (10 μM) 1 μl; reverse primer for V3-V4 amplification (10 μM) 1 μl; 2× Taq Master Mix (Sangon Biotech, B639295, China) 12.5 μl; and 8.5 μl ddH2O (total volume of 25 μl).The mixed reaction system will undergo PCR amplification using the following program in a thermal cycler: denaturation at 94°C for 30 seconds, annealing at 54°C for 30 seconds, extension at 72°C for 1 minute, final extension at 72°C for 10 minutes, and a total of 30 cycles.

#### 2.3.2 Fungal Identification

For fungal identification, the ITS (Internal Transcribed Spacer) region will be amplified using 2× Taq Master Mix (Sangon Biotech, B639295, China). Two universal ITS amplification PCR primers will be utilized: the forward primer ITS1 (5’-TCCGTAGGAACCTGCGG-3’) and the reverse primer ITS4 (5’-CCTCGCTTATTGATATGC-3’). The reaction conditions are as follows: microbial

DNA (10 ng/μl) 2 μl; forward primer for ITS amplification (10 μM) 1 μl; reverse primer for ITS amplification (10 μM) 1 μl; 2 × Taq Master Mix (Sangon Biotech, B639295, China) 12.5 μl; and 8.5 μl ddH2O (total volume of 25 μl).The mixed reaction system will undergo PCR amplification using the following program in a thermal cycler: denaturation at 94°C for 30 seconds, annealing at 54°C for 30 seconds, extension at 72°C for 1 minute, final extension at 72°C for 10 minutes, and a total of 30 cycles.

### 2.4 Construction, quantification, and sequencing of gene library

The samples were sent to Sangon BioTech (Shanghai Shenggong Biotechnology Co., Ltd.) for the construction of gene libraries using universal Illumina adapters and indices. Prior to sequencing, the DNA concentration of each PCR product was determined using the Qubit® 4.0 Green dsDNA Analyzer, and quality control was performed using a bioanalyzer (Agilent 2100, USA). The amplicons in each reaction mixture were pooled in equimolar ratios based on their concentrations.

Following the manufacturer’s instructions, sequencing was carried out using the Illumina MiSeq system (Illumina MiSeq, USA).

### 2.5 Sequence processing, OTU clustering, representative tag alignment, and taxonomic classification

After sequencing, the Illumina short reads were assembled based on overlap using PEAR software (version 0.9.8). The fastq files were processed to generate separate fasta and qual files, and standard methods were employed for analysis. The Usearch software (version 11.0.667) was used to cluster effective tags into Operational Taxonomic Units (OTUs) with a similarity threshold of ≥97%. Chimeric sequences and single OTUs (those with only one read) were removed, and the remaining sequences were sorted into each sample based on the OTUs. The representative sequences of bacterial and fungal OTUs were classified by blasting against the RDP database and the UNITE fungal ITS database, respectively.

### 2.6 Statistical analyses

Alpha diversity indices (including Chao1, Simpson, and Shannon indices) were quantified based on OTU abundance. To assess sample adequacy, observed OTU richness rarefaction curves were constructed, and all alpha diversity indices were calculated using Mothur software (version 3.8.31). OTU rarefaction curves and rank abundance curves were plotted using R software (version 3.6.0). To estimate the diversity of microbial communities in samples, t-tests were used to calculate intra-sample diversity (alpha diversity) for two sample groups, and analysis of variance (ANOVA) was used for multiple group comparisons. Beta diversity can evaluate differences between microbial communities in samples and is often combined with dimension reduction methods such as Principal Coordinates Analysis (PCoA), Non-metric Multidimensional Scaling (NMDS), or Constrained Principal Component Analysis (PCA) for visualization. These analyses were visualized using the R vegan package (version 2.5-6), and distances between samples were displayed in scatter plots. Differential feature analysis was performed using STAMP (version 2.1.3) and LefSe (version 1.1.0) to identify significantly differentially abundant features between groups. The SparCC (version 1.1.0) tool was utilized to calculate correlation coefficients and p-values between communities/OTUs, and the correlation matrix heatmap was generated using the R corrplot package (version 0.84). Network graphs were constructed using the R ggraph package (version 2.0.0).

### 2.7 Relationship between insect eggs and microbial communities

Three experiments were conducted to explore the relationship between insect eggs and microbial communities. The first experiment involved individually selecting three larvae and placing them on blank YPD and LB agar plates to observe microbial growth. The second experiment entailed picking bacteria from plates with visible insect eggs for streaking on agar plates to monitor larval emergence. Lastly, in the third experiment, bacteria were selected and enriched in liquid culture before being spread on agar plates to observe larval presence.

### 2.8 Microscopic observation of larvae with their surrounding flora

The larvae and microbes were stained using the crystal violet staining method to observe their morphological characteristics and sizes under a microscope. The specific steps are as follows: A drop of distilled water was first added to a slide, and a small amount of microbes was picked up with an inoculation loop and mixed with the distilled water. Subsequently, the slide was placed on an alcohol lamp for drying to fix the microbes. Crystal violet staining was applied for 60 seconds, followed by rinsing with water and air-drying for microscopic examination.

### 2.9 Degradation experiments

Screening of cellulose-degrading microorganisms. The formula for carboxymethyl cellulose sodium (CMC-Na) medium used to screen bacteria capable of cellulose degradation is as follows (per liter): CMC Na 15.0g, Na2HPO4·7H2O 12.8g, KH2PO4 3.0g, NaCl 0.5g, NH4Cl 1.0g, agar 15.0g; The formula for carboxymethyl cellulose sodium (CMC-Na) medium used to screen fungi capable of cellulose degradation is as follows (per liter): CMC Na 2.0g, KH2PO4 1.0g, MgSO4 0.5g, (NH4)2SO4 2.0g, NaCl 0.5g, agar 18.0g; No additional carbon source is added to the CMC-Na medium to ensure cellulose is the sole carbon source.

#### 2.9.1 Screening of lignin-degrading microorganisms

The formula for guaiacol broth medium used to screen bacteria capable of lignin degradation is as follows (per liter): 9mmol guaiacol, yeast extract 50mg, Na2HPO4·7H2O 12.8g, KH2PO4 3.0g, NaCl 0.5g, NH4Cl 1.0g, agar 15.0g, with a small amount of yeast extract added to the medium as an auxiliary carbon source component; The formula for guaiacol broth medium used to screen fungi capable of lignin degradation is as follows (per liter): Adding 9mmol guaiacol on YPD medium and using the appearance of a red degradation zone as a screening criterion.

## 3. Results

### 3.1 Microbial morphological analysis under scanning electron microscope

Analysis of microbial morphology on the surface of the mask was conducted using a scanning electron microscope. Various forms of microbes were observed, with the highest presence of observable microbes on the surface M1 (Figure 3a, b, and c), predominantly spherical in shape with occasional rod-shaped structures. Some cells exhibited noticeable concavity (Figure 3a), possibly due to insufficient water supply caused by sampling during the germination process of spores[18], leading to dehydration collapse. Surface M4 (Figure 3f and g) and M5 (Figure 3h) showed a minimal presence of spherical microbes. No microbes with typical morphological characteristics were observed on the surface M3 (Figure 3d and e).The surface and storage environment of M2 are similar to those of M1, M3, M4, and M5. However, during the sample recovery process, the M2 specimen sustained damage. In consideration of preserving cultural heritage and respecting the wishes of the mask collectors, we opted not to conduct a secondary sampling on the M2 mask.

**Figure 3.**
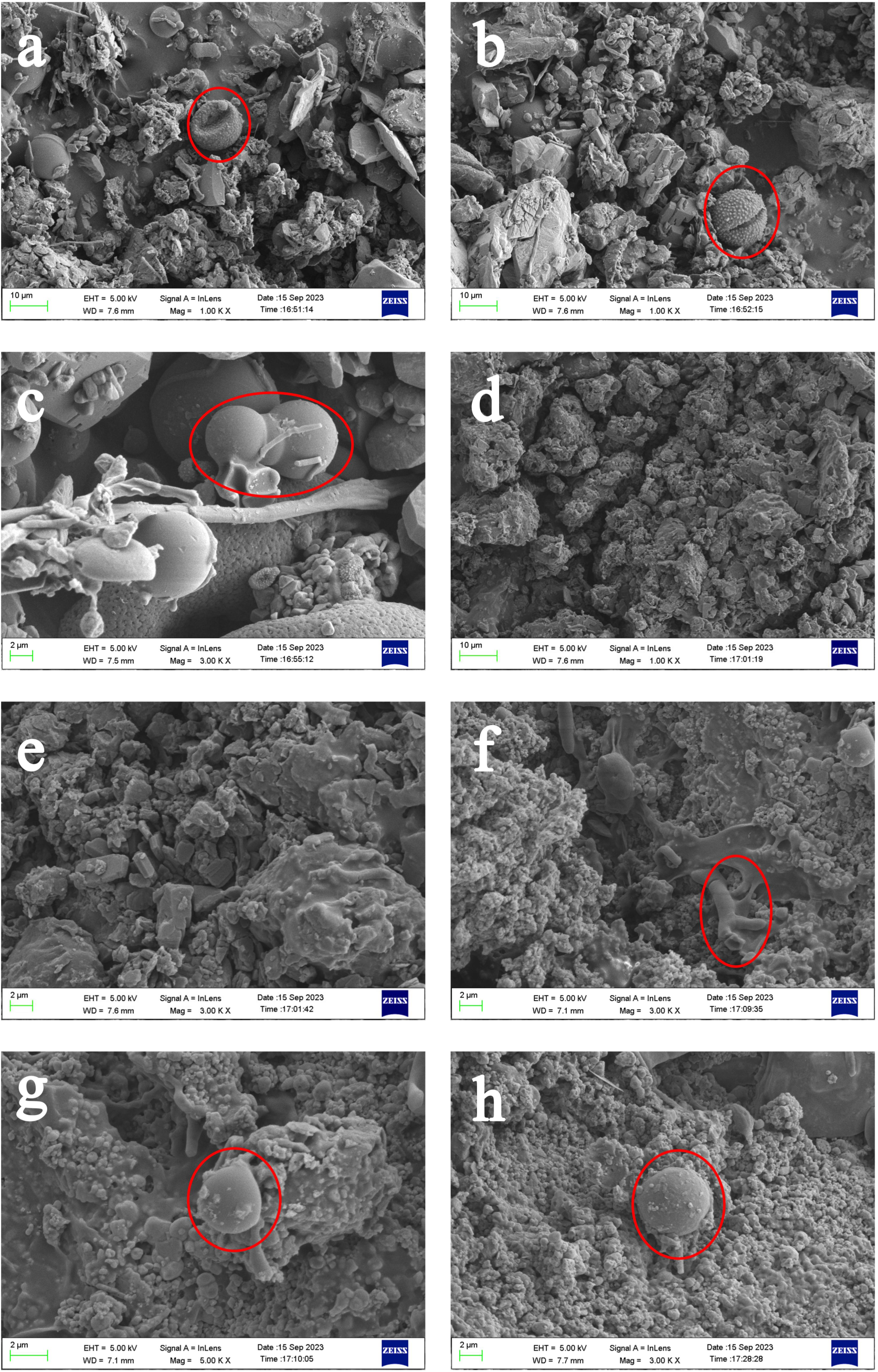
SEM image of the mask sample. (a, b, and c) M1; (d and e) M3;(f and g) M4; (h) M5; (Mag. 1.00-5.00KX), viewing distances 7.1-7.7mm. Distinct colony morphologies can be observed in the scanning electron micrographs. Sample from M2 was damaged, preventing further observation, while no apparent colonies were observed upon examination of the sample from M3.

### 3.2 Analysis of microbial community colonization on mask surfaces based on 16S and 18S amplicon sequencing

Microbial community richness on mask surfaces was evaluated using various indices (Table S1 and S2). Among the samples, amplification of 16S failed for M2-1 and M2-2. Additionally, amplification of 18S was unsuccessful for samples M1-1, M2-2, M3-1, M3-2, and M3-3. The remaining samples exhibited high-quality sequencing results. Bacterial OTU numbers range from 21 to 28, while fungal OTU numbers range from 22 to 82 among the samples. The OTU numbers for each mask are relatively consistent, possibly indicating a weak correlation between OTU numbers and mask storage time. Evaluation of sample biodiversity and microbial richness was performed using Shannon and Simpson indices. The bacterial and fungal Shannon indices for each mask are generally similar (except for M4), with Mask 4 exhibiting the highest bacterial and fungal biodiversity among its sections. Masks 3 and 5 show lower bacterial biodiversity across sections, likely influenced by differences in mask materials affecting microbial colonization. Masks 1 and 2 exhibit locally higher bacterial biodiversity, while Masks 2 and 5 have locally higher fungal biodiversity. The diversity of microbial communities in different sampling sites of some masks was different due to the differences in the age of masks, additional materials (such as horse hair, paint, etc.) and preservation details. Based on the Shannon and Simpson indices, bacterial diversity indices are generally lower, while fungal diversity indices are typically higher across all masks, indicating that fungi are the primary drivers behind the weathering of the masks. Calculations using the coverage index show values close to 1 for all samples, suggesting that our sampling in this study is reasonable and consistent.

Analysis of microbial community composition on masks was conducted at the phylum and genus levels based on sequencing results. At the bacterial phylum level (Figure 4A), Proteobacteria and Firmicutes collectively accounted for over 99% in all masks, being the predominant bacterial phyla on the mask surfaces. Moreover, Proteobacteria predominated over Firmicutes in all masks (except M5-1). At the fungal phylum level (Figure 4B), Ascomycetes constituted over 95% in most of samples (except for an abundance of 24.72% in M1-2). In addition to Ascomycetes, M1-2 exhibited significant representation of Apicomplexa (62.22%). At the bacterial genus level (Figure 4C), Pseudomonas, Paenibacillus (except for <0.5% in M3-1 and <0.2% in M5-2), and Stenotrophomonas (except for <0.1% in M5-1) were notably represented across masks. Specific microbial genera with relatively high abundances uniquely observed in individual masks included Massilia in M1-2 and M5-3 (3.79%, 91.65%), Delftia in M4-1, M4-2, and M4-3 (19.74%, 31.28%, 29.82%), Acinetobacter in M2-3 (50.28%), Exiguobacterium in M1-1, M1-2, and M2-3 (4.72%, 3.22%, 11.01%), Pantoea in M1-2 (5.45%), and Ralstonia (2.40%) and Rhizobium (2.19%) in M5-2.At the fungal genus level (Figure 4D), masks were prominently characterized by Cladosporium, Alternaria (except for absence in M5-2), Aspergillus, Montagnula, and Sarocladium (except for <0.5% in M1-2 and absence in M5-2). Additionally, unique high-abundance fungal genera observed in individual masks included Gregarina in M1-2 and M2-1 (62.22%, 3.22%), norank_Eukaryota in M1-2 (11.93%), Ramularia in M2-1 (6.96%), Fusarium in M4-2 and M5-2 (4.80%, 47.72%), Penicillium in M4-1, M4-2, and M4-3 (1.60%, 5.16%, 2.43%), Kodamaea in M5-2 (3.87%), Acrocalymma and Coniochaeta in M4-2 (3.60%, 3.98%).

**Figure 4.**
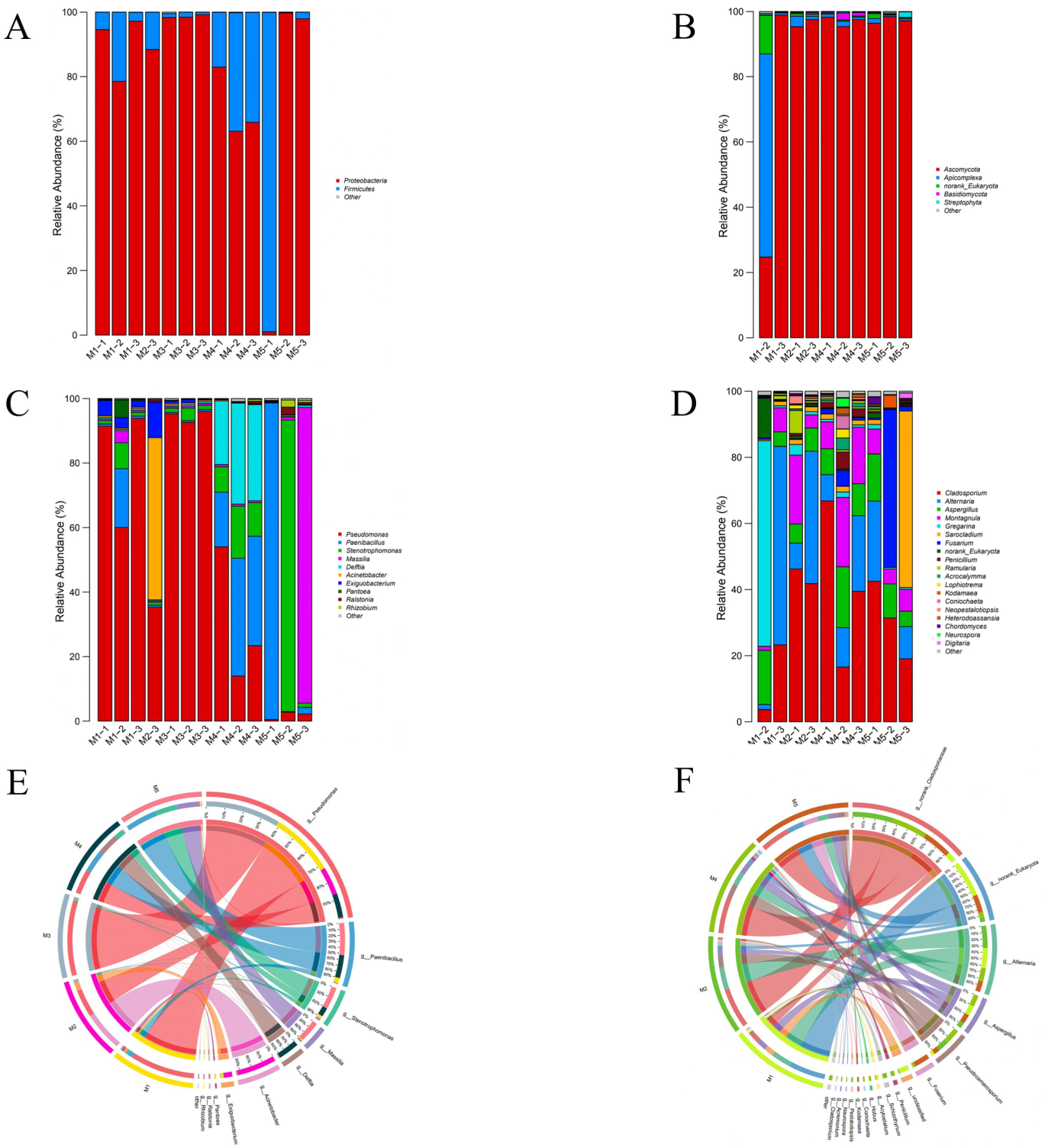
Determination of microbial community diversity in mask samples using amplicon sequencing. (A, B, C, and D) Classification at the phylum and genus levels of microbial communities on the damaged surfaces of five masks under similar climatic conditions based on 16S amplicon and 18S amplicon sequencing. (A) Bacterial phylum; (B) Fungal phylum; (C) Bacterial genus; (D) Fungal genus. (E and F) Co-linearity relationships among mask samples at the genus level based on 16S amplicon and 18S amplicon sequencing. (E) Bacterial genus; (F) Fungal genus.

Co-linearity analysis of dominant microbes on masks. To comprehensively analyze the commonalities and differences in mask microbial communities, samples from five masks (M1, M2, M3, M4, M5) were integrated for further analysis. For bacterial analysis (Figure 4E), Pseudomonas exhibited high abundance across all masks, indicating a commonality among them. Acinetobacter and Exiguobacterium in M2, Massilia in M5, and Paenibacillus and Stenotrophomonas in M4 and M5, highlighting differences among the masks. Regarding fungal analysis (Figure 4F), Cladosporiaceae, Alternaria and Aspergillus displayed relatively high abundances across all masks, suggesting a commonality. Penicillium in M4, and Fusarium in M4 and M5, demonstrating differences among the masks.

### 3.3 Analysis of microbial colonization on the surface of a stage based on 16S and 18S rRNA gene sequencing

Microbial communities colonizing the stage surface were analyzed using 16S and 18S amplicon sequencing. The richness of microbial communities on the stage was assessed using various indices (Table S3 and S4). While amplification of 18S was unsuccessful for sample F, the remaining samples displayed high-quality sequencing results. Bacterial operational taxonomic units (OTUs) ranged from 81 to 137, while fungal OTUs ranged from 83 to 136. Variations in the number of OTUs on the front, sides, and top of the stage indicated a possible correlation between OTU numbers and environmental parameters where artifacts were located. The Shannon index and Simpson index were used to evaluate the biodiversity and microbial richness of the samples. The Shannon index on the front of the stage was relatively consistent and lower compared to the top and sides of the stage. The top of the stage exhibited the highest biodiversity, while the side of the stage showed partial elevation in biodiversity. Calculations based on the coverage index revealed values close to 1 for all samples, indicating adequacy and evenness of the sampling. High diversity indices from the Shannon and Simpson indexes suggested that bacteria and fungi play significant roles in the weathering of the stage from different orientations.

Analysis of microbial community composition on the stage at the phylum and genus levels. At the bacterial phylum level (Figure 5A), Proteobacteria and Firmicutes collectively accounted for over 90% of the stage’s front, sides, top, and skin microbiota (excluding D2). Moreover, Proteobacteria predominated over Firmicutes in all samples. Apart from Firmicutes and Proteobacteria, Actinomycetes were significant in sample D2. Regarding fungal phyla (Figure 5B), Ascomycota and Basidiomycota were abundantly present across all areas of the stage, except for D2 which had less than 1%. Chlorophyta significantly characterized samples C1, C3, D1, D2, and D3, while Ciliophora were notable in samples C2 and D2. At the bacterial genus level (Figure 5C), dominant microbial genera on different parts of the stage and skin exhibited some similarity. All aspects of the stage were significantly represented by Pseudomonas (except for < 1% in Q2), Paenibacillus, and Stenotrophomonas (except for < 0.5% in Q1, Q2, Q3). Specific high-abundance bacterial genera unique to different areas of the stage included Noviherbaspirillum (45.88%) in Q1, Sphingomonas (10.89%) in Q2, Pigmentiphaga (41.92%, 10.15%) and Cellulomonas (3.14%, 3.86%) in Q2 and Q3, Burkholderia-Caballeronia-Paraburkholderia (13.79%, 21.34%) in C1 and C2, Chryseobacterium (28.48%, 4.34%, 5.33%) in D2, D3, and F, and Verticiella (10.25%) in D1.At the fungal genus level (Figure 5D), Cladosporium (except for < 1% in C3) was significant in all directions of the stage and Plenodomus was present in all (except for < 0.5% in C3, D2, D3). Each direction also showed specific genera with relatively high abundances: Colpoda was highly abundant in C2 and D2 (27.33%, 42.52%); Vulcanochloris was highly abundant in D2 and D3 (18.27%, 49.67%); Trichoderma had high abundances in Q3, C1, and C2 (4.00%, 19.51%, 22.50%); Alternaria was highly abundant in Q1, Q2, and D1 (10.29%, 8.73%, 19.30%); Spadicoides was highly abundant in Q1 and C3 (9.73%, 22.99%); Collophorina and Clitopilus were highly abundant in C3 (27.40%, 11.36%); and Rhinocladiella had relatively high abundances in C1 and D3 (8.24%, 9.22%).

**Figure 5.**
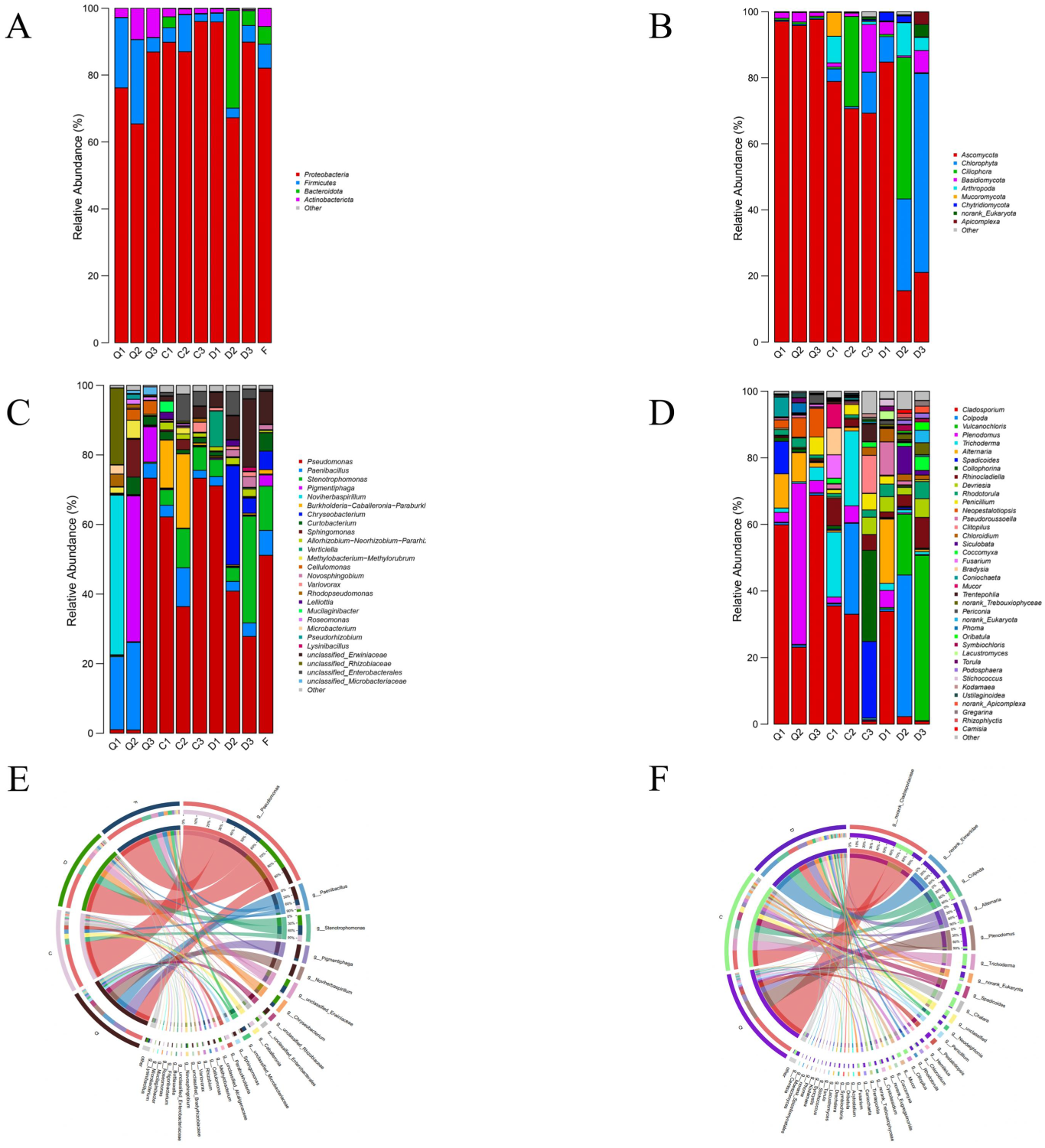
Determination of microbial community diversity in stage samples using amplicon sequencing. (A, B, C, and D) Classification at the phylum and genus levels of microbial communities on the damaged surfaces at the top, front, and side of the stage under similar climatic conditions based on 16S amplicon and 18S amplicon sequencing. (A) Bacterial phylum; (B) Fungal phylum; (C) Bacterial genus; (D) Fungal genus. (E and F) Co-linearity relationships among stage samples at the genus level based on 16S amplicon and 18S amplicon sequencing. (E) Bacterial genus; (F) Fungal genus.

Co-linearity analysis of predominant microbes on the stage was performed. To comprehensively analyze the commonalities and differences across different areas of the stage, samples from the three locations on the stage (Q, C, D) and the skin (F) were integrated for further analysis. In the bacterial analysis (Figure 5E), Pseudomonas, Paenibacillus, and Stenotrophomonas (except for below 0.5% abundance in the front side) were present across all areas of the stage and on the skin, representing the common characteristics observed in all areas of the stage. Pigmentiphaga and Noviherbaspirillum were found at high abundances in the front side (Q), Burkholderia-Caballeronia-Paraburkholderia in the lateral surface (C), and Chryseobacterium in the top (D) and skin (F) areas, indicating the differences observed across different areas of the stage. In the fungal analysis (Figure 5F), Cladosporiaceae and Alternaria were present at relatively high abundances across all areas of the stage, representing the common features observed in all areas of the stage. Plenodomus and Pestalotiopsis were abundant in the front side (Q), Trichoderma and Chalara were prevalent on the lateral surface (C), Eimeriidae were dominant in the top area (D), reflecting the differences observed across different areas of the stage.

### 3.4 Comparison of microbial communities between masks and the stage

Principal component analysis at the OTU level was used to generalize the homogeneity of microbial communities sampled in stage and mask samples. For the 16SRNA, the stage and mask samples are represented in the center of the PCA plot with 95% confidence intervals (Figure S1A, S1B), while the 18SRNA, although the valid samples are not the same as those analyzed by the 16SRNA, also show the same properties (Figure S1C, S1D). The microbial communities between masks, or between stage and different masks are homogeneous. It is reasonable to classify them.

Principal Component Analysis (PCA) of microbial communities between masks and the stage. For bacterial analysis (Figure 6C), Mask 5 (M5-1, M5-2, M5-3) is situated far from the cluster center, indicating a significant difference in species composition compared to other samples. The remaining samples are densely distributed, suggesting a high degree of similarity in bacterial composition between the stage and masks. In the fungal analysis (Figure 6D), M1-2, D2, and D3 are distanced from the cluster center, indicating a substantial difference in species composition compared to other samples. However, the remaining samples are closely clustered, highlighting a high level of similarity in fungal composition between the stage and masks.

**Figure 6.**
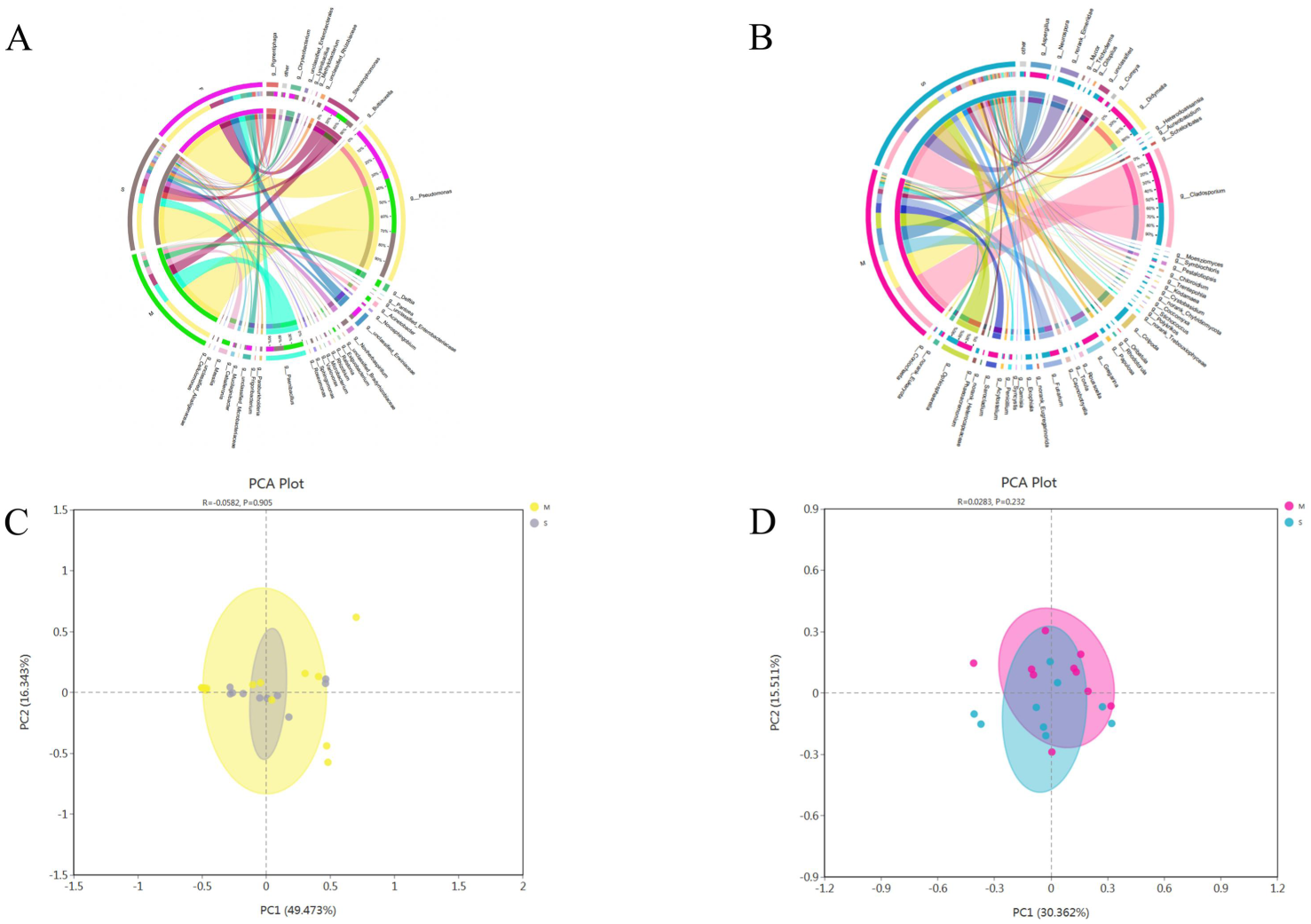
Co-linearity relationships of microbial communities among mask, stage, and human skin samples using amplicon sequencing. (A and B) Co-linearity relationships at the genus level among mask, stage, and human skin samples based on 16S amplicon and 18S amplicon sequencing. (A) Bacterial genus; (B) Fungal genus. (C and D) Analysis of species diversity at the genus level among mask, stage, and human skin samples based on 16S amplicon and 18S amplicon sequencing. (C) Bacterial genus; (D) Fungal genus.

Co-linearity analysis of microbial communities on masks and the stage. We conducted a comprehensive analysis of microbial communities on indoor artifacts represented by masks and outdoor artifacts represented by the stage. For bacterial analysis (Figure 6A), Pseudomonas, Paenibacillus, and Stenotrophomonas are present on masks, the stage, and skin. Pigmentiphaga and Chryseobacterium are found on the stage and skin. Noviherbaspirillum and Burkholderia-Caballeronia-Paraburkholderia exist on the stage. Delftia, Acinetobacter, and Massilia are present on masks. Regarding fungal analysis (Figure 6B), Cladosporium, Ophiosphaerella, and Aspergillus are present on both the stage and masks. Eimeriidae and Colpoda are found on the stage. Gregarina and Sarocladium exist on masks.

### 3.5 Analysis of the relationship between insect eggs and microbial communities

Three larvae were placed on blank LB agar plates and YPD agar plates, and after 7 days, white colonies appeared on both plates. Larvae placed on LB agar showed low activity and no distinct crawling traces (Figure 11Aa). Continuing cultivation for around 20 days on the medium revealed new larvae on the plate (Figure 11Ab), indicating that larvae can carry out normal life activities in the nutritional provided by LB agar and microbial growth environment. Upon magnifying the larvae by 100 times for microscopic observation (Figure 11A and B), their morphology appeared relatively uniform. Subsequently, a small amount of the microbial community around the insect eggs was selected, stained with crystal violet, and magnified 1000 times for microscopic observation (Figure 11Ca and Da). It was observed that there were white granular aggregated substances resembling insect eggs (Figure 11Cb and Db). The insect has been identified as belonging to the Anobiidae family. Its larvae are typically white or cream-colored, with a curved body and distinct head. They feed on wood and are commonly found in furniture, structural timber, and improperly stored wooden artifacts.

### 3.6 Verification of microbial degradation capability based on conventional cultivation methods

Thirteen bacterial strains (Figure 7 and Table S5) and seventeen fungal strains (Figure 8 and Table S6) were isolated from masks and stages using conventional cultivation methods. The ability of 30 strains to degrade cellulose and lignin was evaluated based on CMC-Na agar medium and guaiacol agar medium. Results indicated that three bacterial strains could simultaneously degrade lignin and cellulose (Figure 9a-f), while two bacterial strains could degrade lignin but not cellulose (Figure 9g and h). Additionally, one fungal strain was capable of degrading both cellulose and lignin (Figure 10A and B), whereas twelve fungal strains could degrade cellulose but not lignin (Figure 10a-l).

**Figure 7.**
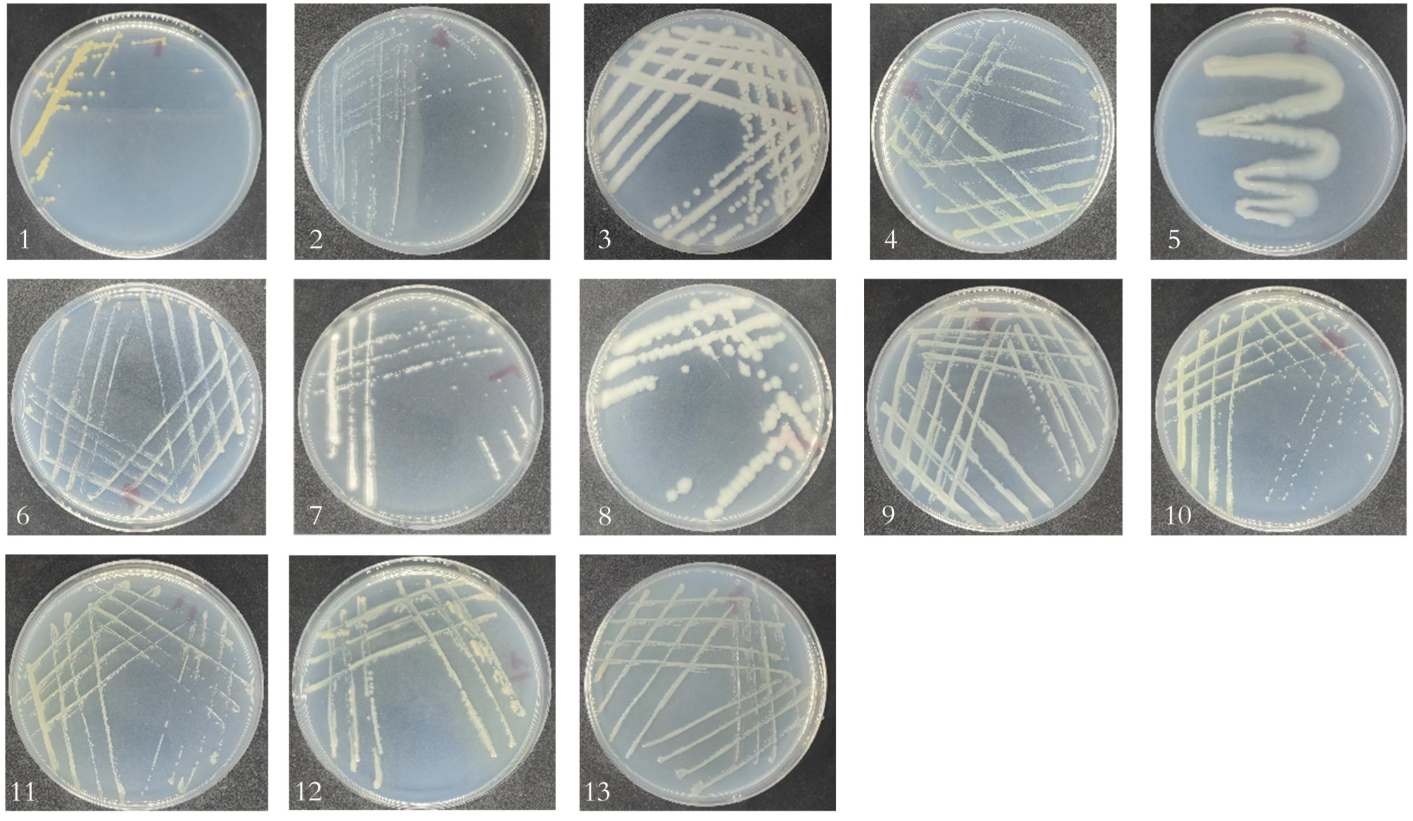
Isolation and purification of bacteria. (1-13) Thirteen bacterial species isolated and identified from microbial samples on masks and stages. Figures 1-13 correspond to B1-B13 in Table 5.

**Figure 8.**
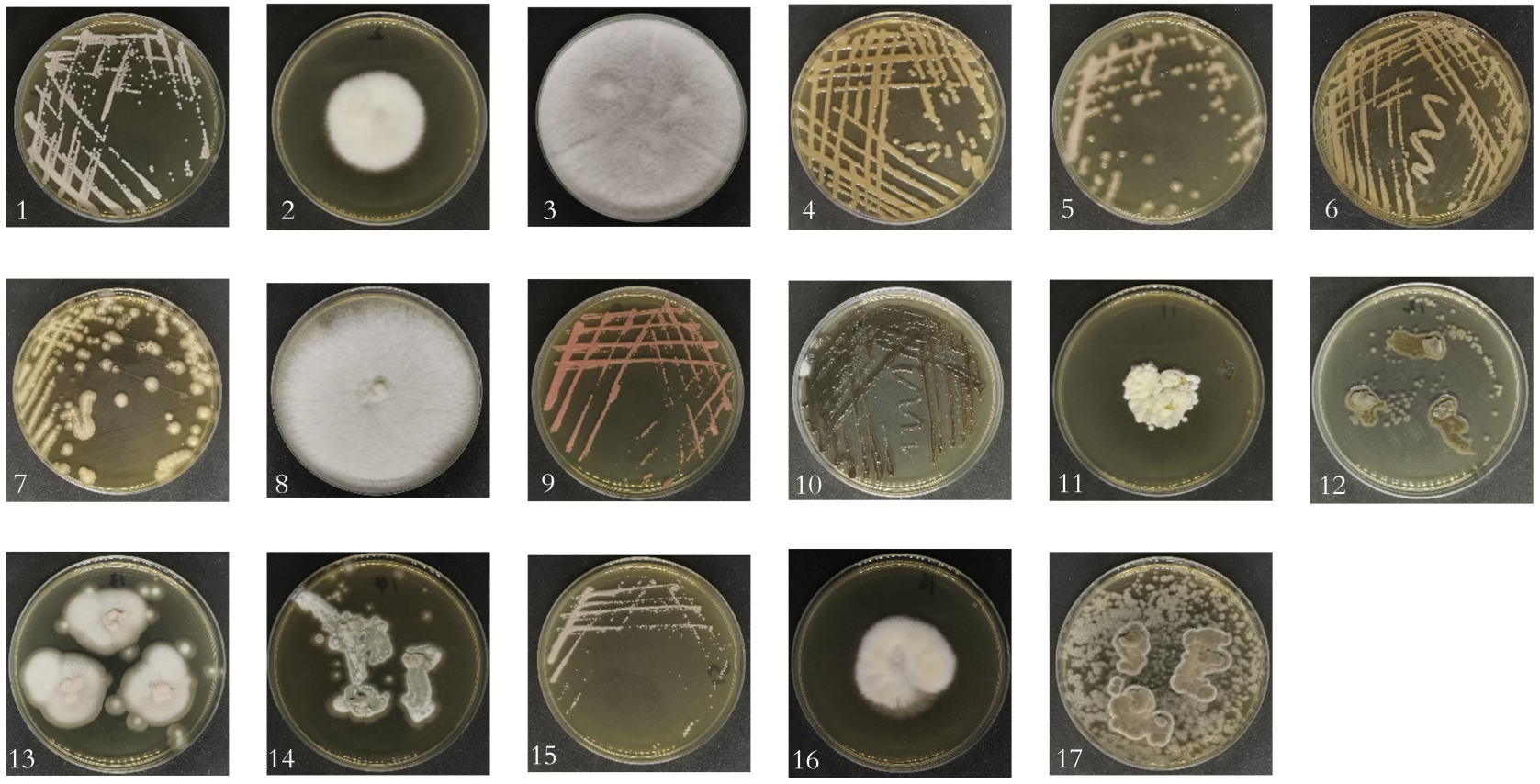
Isolation and purification of fungi. (1-17) Seventeen fungal species isolated and identified from microbial samples on masks and stages. Figures 1-17 correspond to F1-F17 in Table 6.

**Figure 9.**
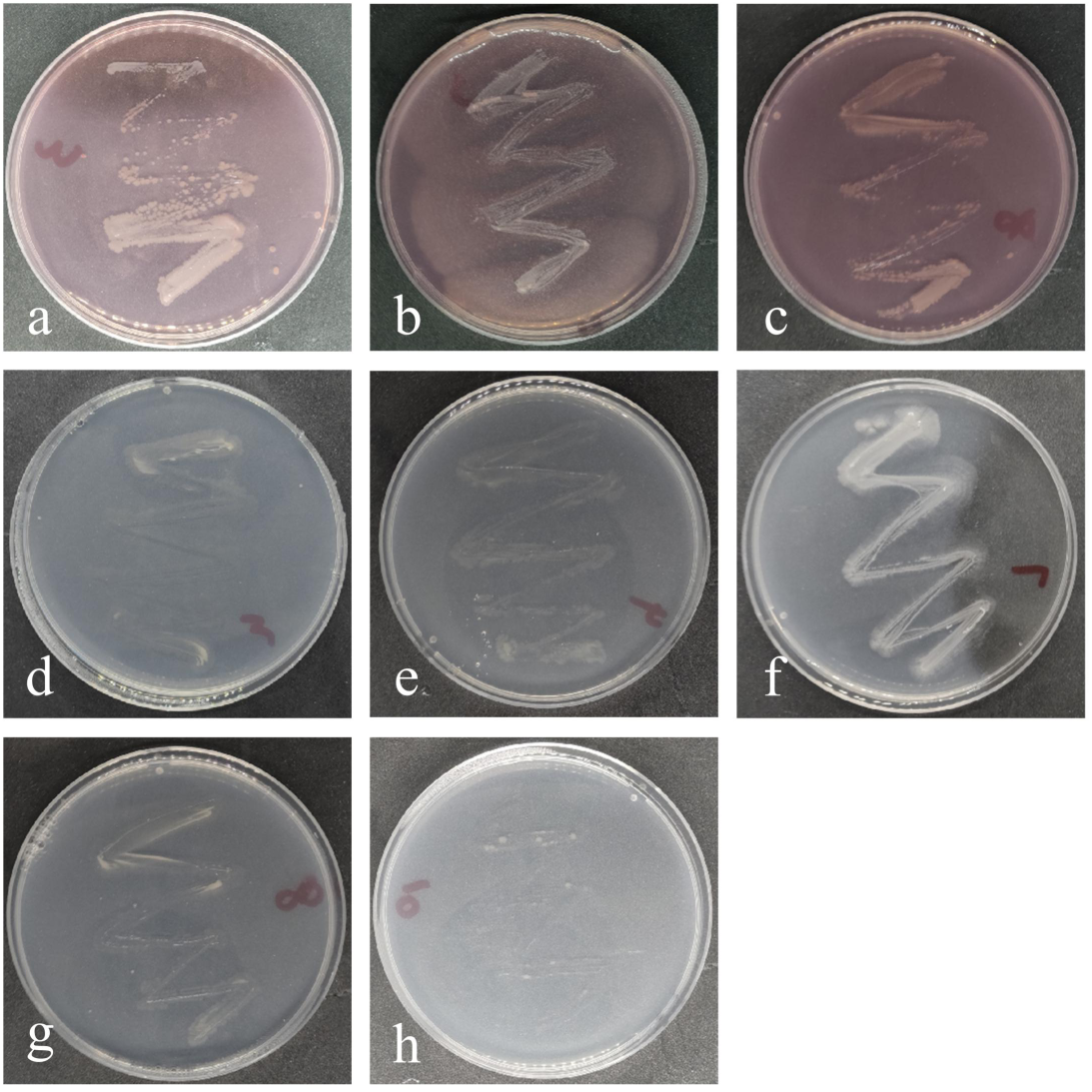
Cellulose and lignin degradation experiments in bacteria. (a, b, and c) Bacteria capable of cellulose degradation; (a) B3, (b) B7, (c) B8. (d, e, f, g, and h) Bacteria capable of lignin degradation; (d) B3, (e) B4, (f) B7, (g) B8, (h) B10.

**Figure 10.**
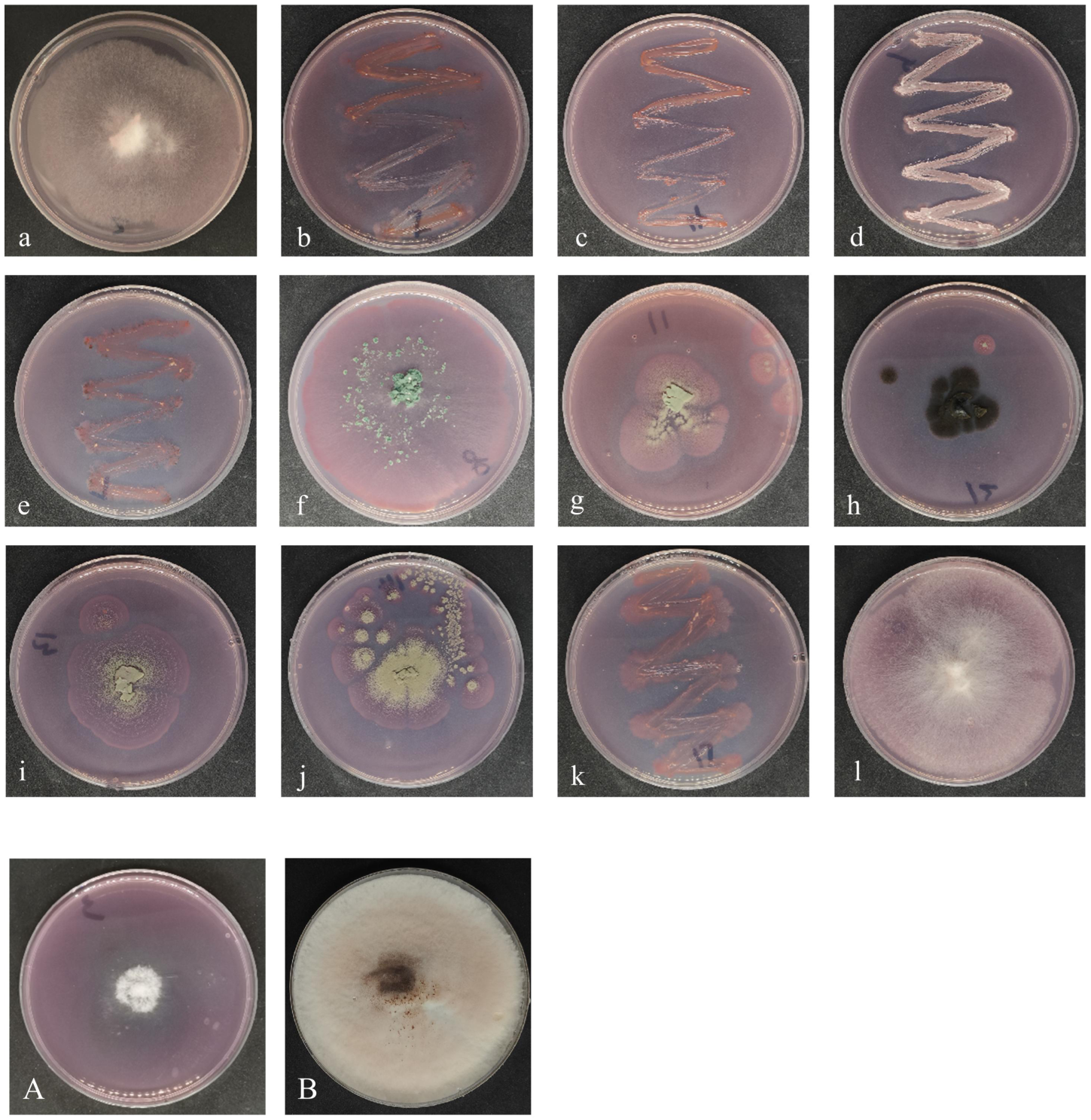
Cellulose and lignin degradation experiments by fungi. (a-l) Cellulose degradation; Figures a-l correspond to B2, B4, B5, B6, B7, B8, B11, B12, B13, B14, B15, B16. (A) Lignin degradation by B3; (B) Cellulose degradation by B3.

**Figure 11.**
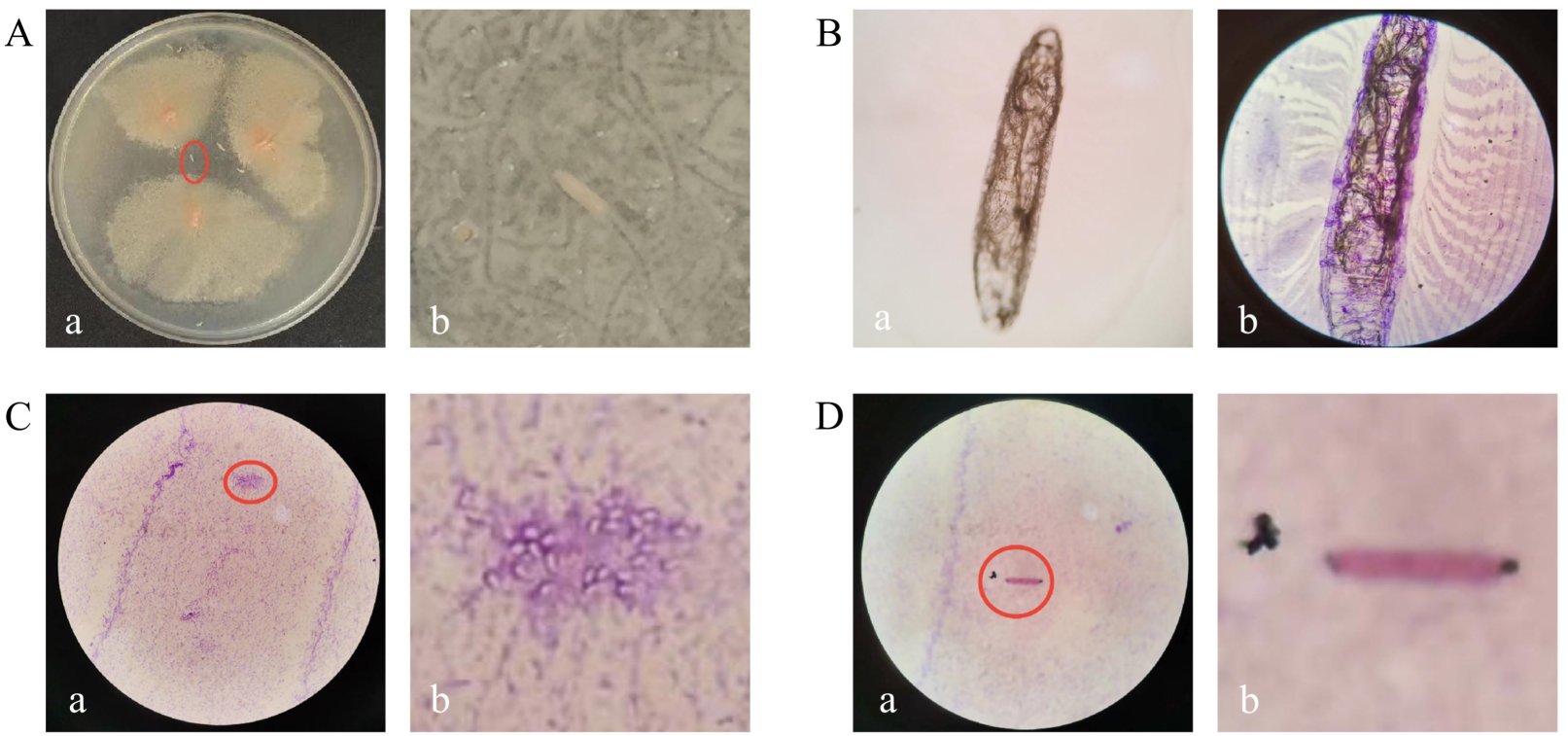
Microscopic observation of worms and colonies. (A)Insects and colonies on LB agar: (a) Overview, (b) Close-up (larva). (B) Insects under microscope at 100x magnification: (a) Unstained, (b) Stained. (C) Bacteria under microscope at 1000x magnification: (a) Overview, (b) Close-up (possible insect eggs).(D) Bacteria under microscope at 1000x magnification: (a) Overview, (b) Close-up (possible larvae).

In all bacteria, Pseudomonas tructae strain, Acinetobacter lwoffii strain, and Bacillus cereus strain are capable of degrading lignin and cellulose, while Stenotrophomonas rhizophila strain and Stenotrophomonas maltophilia strain can degrade lignin. Among all fungi, Rigidoporus vinctus strain can degrade both cellulose and lignin. Additionally, Fusarium oxysporum strain, Hannaella luteola strain, Aureobasidium pullulans strain, Moesziomyces aphidis strain, Aureobasidium leucospermi strain, Trichoderma atroviride strain, Penicillium citreonigrum strain, Cladosporium parahalotolerans strain, Penicillium citrinum strain, Talaromyces verruculosus strain, Sarocladium strictum strain, and Fusarium fujikuroi strain are capable of degrading cellulose.

### 3.7 Bacteria and fungi on the surface of wooden cultural heritage under different climates

We analyzed the bacteria of three climates (Subtropical monsoon climate[19, 20] Temperate broad-leaved forest climate[21], Mediterranean climate[22]) and the fungi of four climates (Subtropical monsoon climate[19, 23], Temperate broad-leaved forest climate[21], Warm temperate monsoon continental climate[24], Boreal ice sheet climate[25]). Specifically, we conducted a detailed analysis of the microorganisms on the surface of wooden artifacts in Guizhou Province (Subtropical monsoon-GZ), Zhejiang Province (Subtropical monsoon-ZJ), and Guangdong Province (Subtropical monsoon-GD), all belonging to the subtropical monsoon climate. Analysis of bacteria at the phylum level (Figure 13A) revealed that Proteobacteria, Actinobacteriota, Firmicutes, and Bacteroidota were common across the three different climates. Additionally, Mediterranean climate exhibited a significantly higher abundance of Cyanobacteria (8.58%). At the genus level (Figure 13C), the shared bacterial genera included Pseudomonas (excluding Temperate broad-leaved forest). Subtropical monsoon-GZ had higher abundances of Paenibacillus (12.62%) and Stenotrophomonas (9.36%), Subtropical monsoon-ZJ exhibited higher abundances of Noviherbaspirillum (15.76%) and Sphingomonas (8.26%), while Subtropical monsoon-GD showed higher abundances of Xanthobacter (5.61%) and Acidovorax (5.55%). Temperate broad-leaved forest displayed a higher abundance of Promicromonospora (8%), whereas Mediterranean climate had higher abundances of Micrococcus (5.99%) and Paracoccus (5.36%) among the bacterial genera. Analyzing the fungal phyla (Figure 13B), common phyla across the different climates included Ascomycota (excluding Warm temperate monsoon continental at 0.28%). Subtropical monsoon-ZJ exhibited a significant abundance of Chlorophyta (41.12%), while Temperate broad-leaved forest showed high abundance of norank_Eukaryota (62.65%) and Warm temperate monsoon continental was characterized by a significant presence of Basidiomycota (99.44%). Looking at the fungal genera (Figure 13D), there were no shared fungal genera among the different climates. Subtropical monsoon-GZ had a high abundance of Cladosporium (27.61%), Subtropical monsoon-ZJ had relatively high abundances of Desmochloris (17.94%) and Phaeosphaeria (14.07%), Subtropical monsoon-GD displayed high abundances of Fusarium (67.65%) and Volutella (14.17%), Temperate broad-leaved forest showed high abundances of norank_Eukaryota (62.65%) and Aspergillus (22.22%), Warm temperate monsoon continental was characterized by a significant presence of Hypochnicium (99.15%), and Boreal ice sheet had relatively high abundances of norank_Helotiales (16.28%) and norank_Chaetothyriales (14.04%).

Based on the PCA plots at the genus level (Figure 13E and F), it can be observed that the distribution of samples under different climates is quite scattered, indicating significant microbial differences on the surfaces of samples from different climatic regions. This suggests that microbial colonization and growth are closely related to environmental conditions such as temperature, humidity, light exposure, and pH.

Even within the Subtropical monsoon climate category, samples from Guizhou (GZ), Zhejiang (ZJ), and Guangdong (GD) in China exhibit substantial microbial diversity on their surfaces. This variability may be attributed to the influence of surrounding environmental factors on microbial colonization. For instance, Guangdong, being a coastal region, experiences high humidity saturation, hot and humid climates, and abundant sunlight. Guizhou, dominated by highland mountainous terrain with well-developed karst ecosystems, exhibits lower temperatures and reduced sunlight exposure. Zhejiang undergoes significant seasonal climate variations, influenced not only by maritime and Southeast Asian monsoons but also by the dual impact of western and eastern air currents, leading to extreme weather phenomena like plum rain and dry spells. Hence, when analyzing the bioerosion status of cultural artifacts across different regions, it is essential to consider not only typical climatic conditions but also specific geographical features.

## 4. Discussion

The main focus of this project lies in investigating the microbial weathering processes on wooden masks and wooden ancient opera stages. It involves identifying the core microbial communities shared by indoor and outdoor artifacts in karst landform environments, validating purified microbes’ capabilities in degrading lignin and fibers, exploring the relationship between insect eggs and microbes, and summarizing microbial differences under varying climatic conditions.

Analysis based on species composition and abundance bar graphs at the 16S and 18S taxonomic levels (Figure 4A-D) and OTU-level PCA analysis (Figure S1A and S1C) showed significant differences among bacterial species on different masks, while fungi demonstrated greater commonalities. This suggests that the colonization and growth of different bacteria on masks may be closely linked to the preservation time and material composition, whereas the impact of different fungi appears to be less pronounced. In indoor settings with stable temperature and humidity conditions, a predominant core microbial community contributing to mask weathering includes bacteria such as Pseudomonas, along with fungi like Cladosporium, Alternaria, Aspergillus. These microbial strains are less influenced by the duration of artifact preservation or the material composition, serving as the primary genera causing microbial weathering of wooden artifacts indoors.

Analysis based on species composition and abundance bar graphs at the 16S and 18S taxonomic levels (Figure 5A-D) and OTU-level PCA analysis (Figure S1B and S1D) revealed considerable differences among fungal species across various positions on opera stages, while bacteria shared more similarities. This indicates that the colonization and growth of different fungi on opera stages at different positions may be closely associated with factors like light exposure, ventilation, wood types, whereas the relationship for bacteria seems less significant. In outdoor exposed environments with fluctuating temperature and humidity, a fixed core microbial community responsible for opera stage weathering includes bacteria such as Pseudomonas, Paenibacillus, Stenotrophomonas, and fungi like Cladosporium, Alternaria. These microorganisms exhibit strong colonization and growth abilities, thriving widely across different positions and wood types on opera stages outdoors, representing the key genera causing microbial weathering of wooden artifacts in outdoor settings.

Comparing the Shannon and Simpson indices (Table S1-S4), it is evident that the bacterial and fungal diversity indices of the stage are generally higher than those of the mask. This suggests that outdoor microbial species are richer compared to indoor environments, and microbial weathering phenomena are more severe outdoors. Analysis through co-linearity figures (Figure 6A and B) and PCA analysis (Figure 6C and D) indicates that there is minimal difference in species composition of bacteria between masks and stages at the genus level. In contrast, there is significant variation in the species composition of fungi between masks and stages. This suggests that fungal diversity on cultural heritage changes significantly between indoor and outdoor environments, whereas bacterial diversity changes minimally. Hence, fungi colonization is more susceptible to environmental changes. Under karst topography and climatic conditions, both indoor and outdoor cultural heritage’ surfaces harbor a core microbial community responsible for weathering. This core group includes bacteria represented by Pseudomonas, Paenibacillus, and Stenotrophomonas, as well as fungi represented by Cladosporium, Alternaria, and Aspergillus. These microbes can colonize and grow in both indoor and outdoor environments, making them the primary genera responsible for microbial weathering of wooden cultural heritage indoors and outdoors.

Specific analysis of the dominant bacterial genera on the surfaces of indoor and outdoor cultural heritage. Pseudomonas is predominantly an aerobic bacterium that relies on strict respiratory metabolism. However, it allows anaerobic growth when nitrate serves as an electron acceptor[26]. Consequently, Pseudomonas thrives in well-ventilated areas where oxygen is abundant, while its growth in anaerobic environments may be associated with nitrogen cycling. Pseudomonas can become a major microorganism indoors and outdoors, possibly due to its ability to colonize and grow normally under both aerobic and anaerobic conditions. Moreover, Pseudomonas is commonly found on human skin[27], as confirmed by skin samples from this study, indicating a relatively higher abundance of Pseudomonas on the skin compared to other bacterial species. Therefore, human activities may serve as a medium for the spread of this bacterium, influencing the colonization and growth of Pseudomonas on the surfaces of indoor and outdoor cultural heritage artifacts. Paenibacillus is primarily an anaerobic bacterium that produces acids[28]. These secretions may cause certain damage to the artistic value of artifacts. Research suggests that Paenibacillus is widely present in different soils and can affect plant growth[29]. Thus, Paenibacillus is more prevalent in outdoor environments compared to indoor settings. It can trigger systemic resistance or secrete substances to kill pathogens and possesses genes related to nitrogen fixation, phosphorus assimilation, and iron metabolism[30]. This indicates that Paenibacillus can provide nutritional substances through nitrogen fixation and phosphorus assimilation for itself and surrounding plants, giving it a significant advantage in nutrient competition over other bacterial species and leading to its widespread colonization on indoor and outdoor cultural heritage surfaces. Biofilm is a high-molecular matrix composed of polysaccharides, proteins, lipids, nucleic acids, and bacteria with low metabolic activity. It can rapidly mature and settle on new surfaces within 24 hours[31–33]. Biofilm formation is a primary mechanism through which microorganisms damage the surfaces of artifacts. Studies have shown that Stenotrophomonas exhibits a strong ability to form biofilms[34], and is commonly found in aquatic environments, plants, and soils. Moist surfaces are conducive to bacterial proliferation and growth, and Stenotrophomonas can also colonize plastic or metal surfaces. Therefore, failure to periodically dry artifact surfaces may result in the adhesion and extensive growth of Stenotrophomonas, causing irreversible damage to artifacts through attachment.

Specific analysis of the dominant fungal genera on the surfaces of indoor and outdoor cultural heritage. Cladosporium is commonly found in the dead stems and leaves of woody plants, making this type of fungi more likely to colonize and grow on the surfaces of outdoor cultural heritage near trees. For instance, in front of the ancient stage sampled in this study, there are numerous shrubs nearby providing suitable conditions for Cladosporium colonization. Research indicates that such fungi can be easily isolated from air, fabrics, paints, and other organic materials[35, 36]. Moreover, their conidia are very small and highly capable of long-distance dispersion in the air[36, 37]. Hence, Cladosporium on the front side of the stage is likely to extensively colonize various areas of the stage through this mode of spread. Cladosporium can parasitize and proliferate within other fungi, with their natural metabolites exhibiting antibacterial properties[38]. This characteristic may pose a threat to the host and other fungi, allowing Cladosporium to gain dominance in population competition and become the predominant fungal genus on both indoor and outdoor cultural heritage.

Alternaria can thrive in various substrates[39]. Their spores disperse in dry air[40, 41], while high humidity conditions favor their colonization, growth, and mycotoxin production[42]. Therefore, Alternaria spores might spread widely during relatively dry seasons and settle and grow on artifact surfaces during high humidity in summer.

Alternaria has a broad temperature tolerance range, with an optimal temperature of 20 ℃ [40], similar to our indoor sampling environment. Suitable temperature and humidity conditions may be key factors contributing to Alternaria becoming a predominant microbial genus on mask surfaces. Additionally, strains of this genus produce black pigments[40], affecting the aesthetics of artifacts. Aspergillus is a filamentous fungus associated with human lung diseases[43], hence its environmental spread may be related to human activities. Aspergillus has a wide adaptability range and can commonly grow in deserts, forests, wetlands, and cultivated soils worldwide[44]. Aspergillus secretes pigments, proteolytic enzymes, and mycotoxins[45]. Not only can it degrade cellulose, but also its mycelial growth exerts mechanical pressure on substrates, causing physical damage to artifacts[18]. The strong adaptability of Aspergillus enables it to colonize and grow on artifact surfaces, leading to discoloration of surface materials and increased fragility. Furthermore, Airborne fungal spores or metabolic byproducts may contaminate indoor environments[46].

Previous researchers have been focusing on identifying the microorganisms present on artifacts without paying much attention to where these microorganisms come from. During our purification process, we discovered white larvae which is identified as Anobiidae family in the culture medium. Under inadequate sealing conditions, these larvae carrying bacteria would crawl onto other culture media and contaminate other bacterial strains. Therefore, based on this observation, we speculated that these larvae may serve as a means of transmission and initiated related experiments to verify this hypothesis. We identified white colonies appearing after placing larvae in LB and YPD culture media and created bubble charts (Figure 12A and B). It was observed that microorganisms related to the larvae, such as Pseudomonas, Stenotrophomonas, as well as fungal genera like Cladosporium, Alternaria, Aspergillus, and Fusarium, overlapped with the core microorganisms present indoors and outdoors in Anshun. Additionally, an unidentified eukaryotic organism was identified, comprising 3.3035% of the total, but due to a lack of demonstrable functionality, it was not discussed.Further analysis of the core microorganisms related to the larvae revealed their shared capability for cellulose or lignin degradation. Moreover, Stenotrophomonas exhibited biofilm formation ability, while Cladosporium has the capacity to produce mold toxins to compete against other fungi, enhancing its own competitive advantage, and Alternaria can produce melanin to contaminate the surfaces of cultural relics. These microorganisms form a communication network among the stage, masks, and larvae, potentially adhering to the surface of the larvae, with the movement of the larvae expanding the spread of these microbial species.During the observation of white colonies, an image resembling a larva egg was detected (Figure 11Cb), suggesting that these microbial strains could potentially carry larva eggs during normal transmission and inoculation processes, leading to bidirectional transmission(Figure S2). Moreover, these larvae eggs could reproduce and thrive normally on the culture medium without being influenced by the microbial environment. Based on the experimental results mentioned above, these larvae eggs act as crucial intermediaries facilitating the transmission and inoculation of different microbial communities. The larvae can carry these microbial communities for their normal life activities, thereby enhancing interactions between diverse microbial communities and expanding the colonization growth range of microorganisms, ultimately causing more significant damage to artifacts. Preservation conditions for different artifacts vary due to practical considerations, making it challenging to implement uniform protection measures.

**Figure 12.**
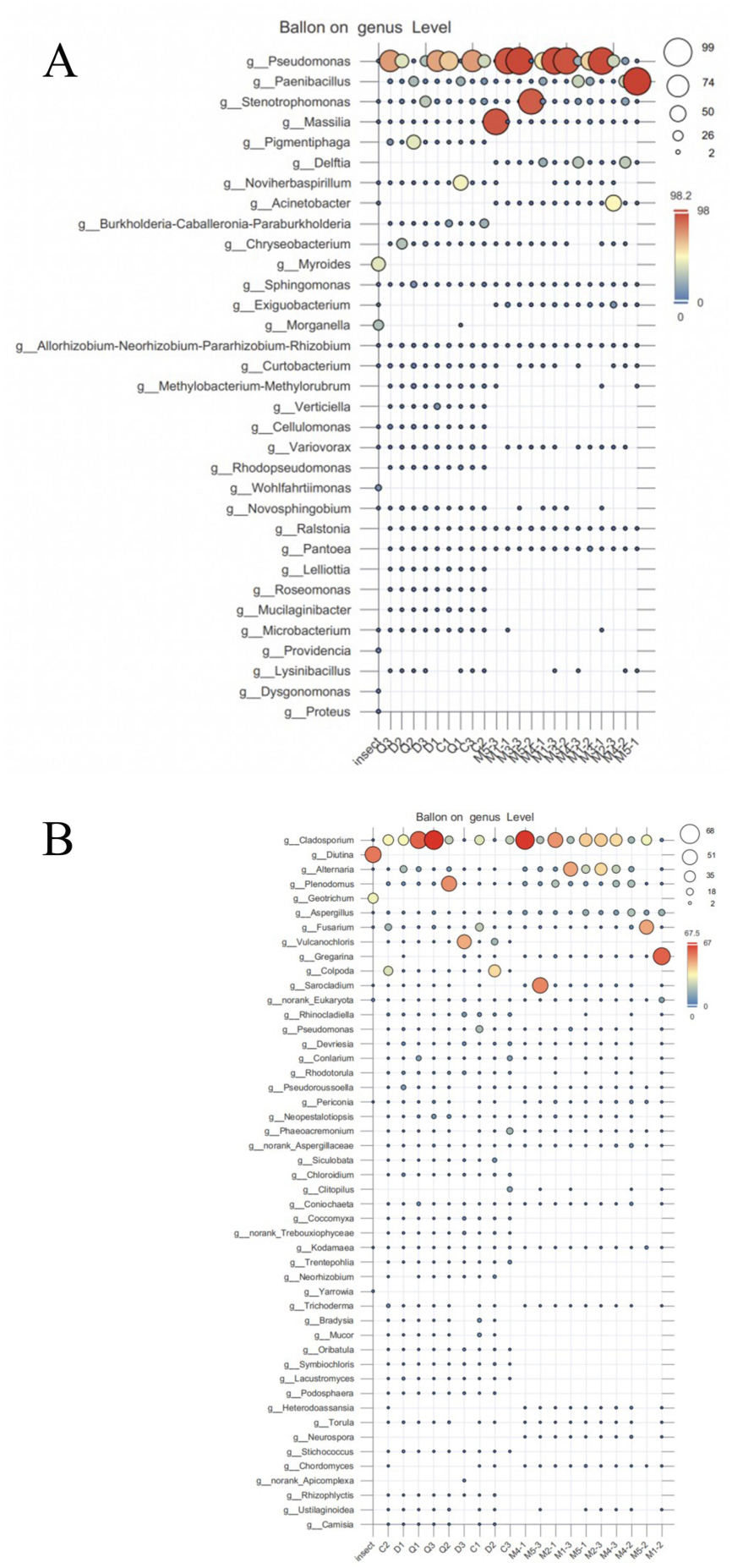
Bubble plots of microorganisms on insect surfaces and masks, theatre sampling samples. Genus-level bubble chart. (A) Bacterial genera; (B) Fungal genera. The larger the bubble, the higher the abundance of the species on the sample.

**Figure 13.**
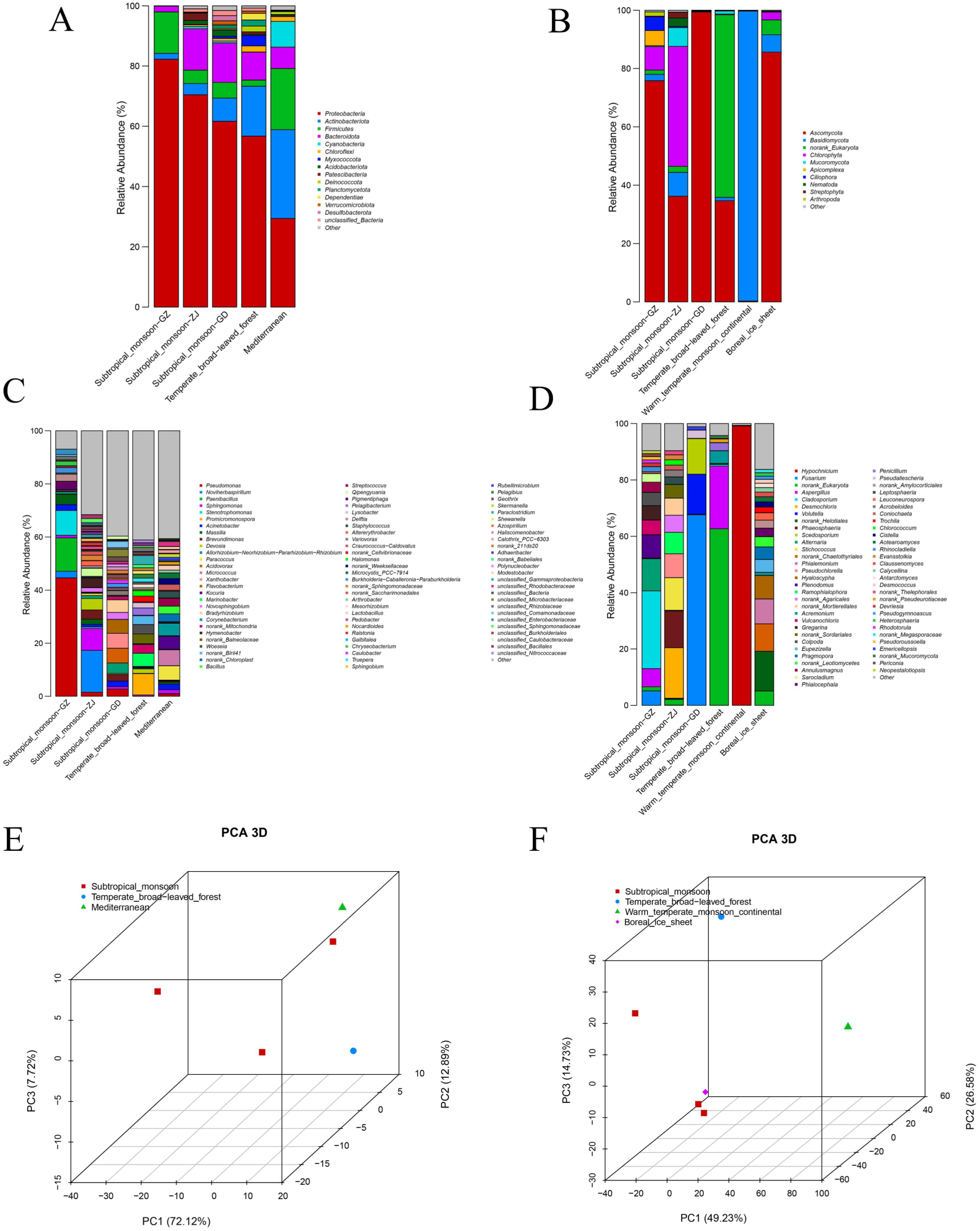
Diversity of microbial communities in woody artefacts from different climates determined using amplicon sequencing. (A, B, C, and D) Genus-level classification and species-level classification of microbial communities on the surfaces of wooden artifacts from different climates based on 16S amplicon and 18S amplicon sequencing.(A) Bacterial phyla; (B) Fungal phyla; (C) Bacterial genera; (D) Fungal genera. (E and F) Differential analysis of species at the genus level among samples of wooden artifacts from different climates based on 16S amplicon and 18S amplicon sequencing.(E) Bacterial genera; (F) Fungal genera.

Wooden artifacts like masks and stages are highly susceptible to degradation by microorganisms expressing wood cellulose-degrading enzymes due to the inherent nature of the material. Microbes capable of degrading cellulose utilize it as a carbon source, breaking it down to glucose for sustenance, leading to their growth and reproduction on artifact surfaces. Cellulose typically exists in crystalline form, with some disorganized cellulose chains forming amorphous cellulose, which is more easily degraded by enzymes[47]. Cellulose degradation primarily involves cellulase enzymes (endoglucanases, exoglucanases, and β-glucosidases), with β-glucosidase being the key enzyme hydrolyzing fiber sugars into glucose[48], releasing pigments or weak acids causing rust spots or discoloration on paper surfaces. Lignin degradation requires oxygen-dependent oxidation processes, primarily mediated by three enzymes: lignin peroxidase (LiP), manganese peroxidase (MnP), and laccase[6]. Among these, laccase is a metal-containing oxidative enzyme capable of oxidizing various organic and inorganic substrates[49].

In degradation experiments, 5 bacterial strains and 12 fungal strains exhibited cellulose-degrading activity, while 2 bacterial strains and 1 fungal strain displayed lignin-degrading activity. Studies have shown that Pseudomonas is prevalent in most petroleum-contaminated sites, capable of metabolizing complex hydrocarbons under aerobic conditions using monooxygenase or dioxygenase to degrade oil components, this suggests that they have the ability to metabolize complex hydrocarbons[50, 51].

Stenotrophomonas secretes a variety of enzymes involved in the metabolism of polycyclic aromatic hydrocarbons (PAHs) and has diverse metabolic pathways[52], this could be the reason why it is able to degrade lignin. Fusarium, a member of Ascomycota, expresses a full set of cellulose-degrading enzymes, predominantly high levels of β-glucosidase[53], also producing mycotoxins[54, 55]. Moreover, Fusarium is adsorbent[56], and this property may make it more firmly adsorbed on the surface of artifacts, in addition, it can cause rust and sooty mold[57, 58], so it can not only make wooden artifacts brittle by degrading cellulose, but also may discolor the surface of artifacts. Penicillium is known to possess the function of decomposing organic matter[59], and it has been confirmed to exhibit high cellulose degradation activity[46]. When cellulose is present near the mycelium of Penicillium, it allows cellulases to hydrolyze as inducers. Upon the entry of inducers such as cellulose oligosaccharides into the cell, the activation of proteins and activation elements triggers the transcription of cellulase genes, subsequently degrading cellulose and releasing a large amount of glucose[60]. Trichoderma possesses a powerful cellulase expression system, renowned for its abundant production of cellulases and hemicellulases[61]. Additionally, Trichoderma exhibits faster growth rates compared to other filamentous fungi[62]. Enzymes secreted by Trichoderma related to cellulose hydrolysis, such as β-glucosidase (BGL) and xylanase, have garnered significant attention from the industrial sector[63, 64],demonstrating a remarkable ability to degrade cellulose. It is worth mentioning that several filamentous fungi like Penicillium, Trichoderma, and Aspergillus can produce synergistic cellulases that collectively act on the efficient decomposition of cellulose[65]. Moreover, these filamentous fungi are commonly found at high abundance in sampled specimens, so the interactions among them can lead to profound damage to the internal fiber structure of artifacts. In this study, it was also demonstrated that Aureobasidium pullulans and Sarocladium strictum can degrade cellulose. Both of these fungi exhibit halotolerance, adapting well to saline-alkaline environments[66, 67] Halophilic fungi are xerophilic organisms, and their presence is associated with indoor dust and the coexistence of other xerophilic fungi such as Penicillium[18]. Aureobasidium pullulans is commonly found on plant leaves[68], and wood-degrading enzymes such as xylanase[69] and β-glucosidase[70] have been identified in its enzymatic system. Research has observed under scanning electron microscopy that the mycelium of Sarocladium strictum attaches to the leaf surface and forms a dense network, subsequently invading leaf tissues by secreting cellulase to degrade cellulose entirely[71]. Hence, this fungus may erode cultural artifacts in a similar manner. Rigidoporus vinctus exhibited significant lignin degradation capability in this study. It is a white rot fungus causing tissue discoloration and death in trees, leading to the development of white rot on artifact surfaces, affecting their aesthetics. R. vinctus belongs to the MnP-Lac class of degrading fungi, where the enzymes MnP and Lac collaboratively participate in the "delignification" process, degrading lignin by opening the phenyl ring[72–75]. Additionally, R. vinctus degrades cellulose similarly to other fungi, breaking down cellulose into glucose through the synergistic action of multi-component enzyme complexes[72]. Rhodotorula colonies appear pink, with carotenoids as their secondary metabolites[76]. Exophiala colonies are black, capable of producing abundant melanin through the 1,8-dihydroxynaphthalene (DHN) melanin pathway, embedding it into the cell wall[77]. While these fungi might not degrade cellulose, carotenoids and melanin serve as natural colorants[78] that can cause discoloration on artifact surfaces.

Analyzing dominant microorganisms in different climates, Warm temperate monsoon continental regions showed significant representation of Basidiomycota (99.44%). Basidiomycota typically thrive in dry and cool environments[79], and fungi belonging to Basidiomycota demonstrate significant tolerance to high temperatures and severe cold.[80]. This adaptation aligns well with the hot summers and cold, dry winters characteristic of Warm temperate monsoon continental regions. In Temperate broad-leaved forests, Aspergillus (22.22%) displayed higher abundance. Aspergillus survival in diverse environments and geographical habitats is linked to its metabolic diversity, high reproductive capacity, and competitive edge in nature. It also possesses xerophilic characteristics, thriving poorly below 12°C[81]. The dry winter conditions and temperature fluctuations between 13-23°C in Temperate broad-leaved forests create ideal growth conditions for Aspergillus. Subtropical _ monsoon-GD harbors a high abundance of Fusarium (67.65%), which thrives in humid environments and grows well at temperatures around 0-37°C[82]. Continuous exposure to light is more conducive to its growth[83], which aligns with the climatic characteristics of Guangdong Province, known for its high humidity, heat, and abundant sunlight. Thus, the temperature and humidity preferences of microbial species for growth and reproduction are closely related to different climates. Subtropical _ monsoon-GZ exhibits a higher abundance of Cladosporium (27.61%), which demonstrates various ecological adaptations such as endophytic, plant pathogenic, and saprophytic modes[84]. Due to the diverse plant species in Guizhou Province, investigations have revealed richness in both the quantity and diversity of Cladosporium[37]. This indicates that the relationship between the nutrient pattern and the surrounding environment will affect the colonization range and growth ability of the strain. Boreal ice sheet harbors a higher abundance of norank Helotiales (16.28%) and norank Chaetothyriales (14.04%). Helotiales comprise two types, characterized by either "distinctive morphology and strict fungal structure" or "diffuse or indefinite morphology" in substrates, both considered overwintering structures capable of surviving at low temperatures[85]. Helotiales also express stress-resistant genes, secrete CAZymes, and possess strong competitiveness[86]. Norank Chaetothyriales (14.04%) were generally found in habitats under extreme harsh conditions[87], and fungi belonging to Chaetothyriales are black in color. Melanin has multiple functions to enhance survival under extreme conditions[88]. Additionally, these fungi exhibit oligotrophic characteristics [89]. The combination of these features enables these fungi to survive in the extreme climatic conditions of the Boreal ice sheet. This suggests a correlation between microbial growth under certain extreme conditions and their structural characteristics, expression of specific genes, and nutritional requirements.

## 5. Conclusion

On May 31, 2011, the 22nd meeting of the Standing Committee of the 11th People’s Congress of Guizhou Province approved the promulgation of the "Regulations on the Protection of Tunbao Cultural Heritage in Anshun, Guizhou Province," which would come into effect on August 1, 2011. This regulation provided a legal basis for the protection of Tunbao culture.This research aims to explore the damage mechanisms of Tunbao local opera masks and theatres during microbial weathering processes and how effective protection and restoration can be carried out through scientific methods. By analyzing the microbial communities on Tunbao local opera masks and theatres and considering environmental factors, we will propose targeted prevention and control strategies to offer new perspectives and methods for the protection of Tunbao cultural heritage.Through this study, we hope to contribute to the inheritance and development of Tunbao culture, allowing this unique cultural phenomenon to shine brightly in the new era.

To summarize, this project employed sampling scanning electron microscopy to identify different forms of microbial colonies on the surfaces of artifacts. By using traditional culture methods and high-throughput sequencing, the microbial weathering conditions on the surfaces of five masks representing indoor artifacts and an ancient stage representing outdoor artifacts were analyzed. Pseudomonas, Paenibacillus, Stenotrophomonas, Cladosporium, Alternaria, and Aspergillus were identified as core microorganisms common to indoor and outdoor environments. We found that the colonization of Paenibacillus and Cladosporium is closely related to the vegetation diversity in Anshun region, while the colonization of Stenotrophomonas and Alternaria is somewhat correlated with the temperature and humidity of this area. The dissemination of Pseudomonas and Aspergillus is influenced by human activities. Isolation and identification of samples, along with exploration of their cellulose and lignin degradation capability, led to the characterization of their competitive advantages in terms of physiological characteristics and degradation abilities. It was observed that the colonization of dominant microorganisms on artifact surfaces was correlated with spore dispersal capability, enzyme secretion activity, and types of secondary metabolites (such as fungal toxins). Through microscopic observation of the cultured insect eggs and microbial community, it was identified that the insect eggs belong to the Anobiidae family. Analyzing the correlation between this microbial community and surface microorganisms of various samples confirmed that these eggs serve as a medium to spread microorganisms and expand their colonization range. By inhibiting the transmission pathway of microorganisms through suppressing insect eggs, a new approach for the unified protection of wooden artifacts in Anshun area has been proposed.By analyzing the differences in surface microorganisms on wooden cultural heritage under different climates and the characteristics of dominant microorganisms, it was confirmed that microbial growth and reproduction under varied climatic conditions are correlated with the surrounding environmental climate, vegetation conditions, as well as specific structural features and unique gene expressions of the microorganisms themselves. However, research on surface microbiota of wood artifacts in different climates is still insufficient, thus limiting the depth of analysis in this regard.

## Acknowledgements

We thank all tunbao culture heritage managers for their help in collecting microbiological samples of stage and masks.

## Author contributions

J.R.L and J.L conceived the experiments. J.R.L, P.A and C.G collected the microbiological samples. P.A and X.L performed the experiments. P.A analyzed the data. P.A, X.L and C.G wrote the paper. J.R.L and J.L performed the review and editing of the original draft.

## Data availability

The amplicon sequences and shotgun metagenomics reported in this article have been deposited in the NCBI BioProject (accession nos. PRJNA1146571).

