## Supplemental table and figure for "Microbial Weathering Analysis of Anshun Tunbao Artifacts"

### Supplementary material

**Table S1. Bacterial diversity index analysis table for mask samples**

| Sample | Number | OTUs | Shannon | Chao | Ace | Simpson | Shannoneven | Coverage |
|---|---|---|---|---|---|---|---|---|
| M1-1 | 47401.0 | 25.0 | 0.57 | 35.0 | 37.83 | 0.79 | 0.18 | 1.00 |
| M1-2 | 49944.0 | 27.0 | 1.65 | 29.5 | 38.44 | 0.33 | 0.50 | 1.00 |
| M1-3 | 49951.0 | 25.0 | 0.63 | 28.0 | 29.61 | 0.76 | 0.20 | 1.00 |
| M2-3 | 50193.0 | 23.0 | 1.21 | 24.5 | 25.74 | 0.38 | 0.39 | 1.00 |
| M3-1 | 51458.0 | 23.0 | 0.45 | 24.0 | 24.23 | 0.85 | 0.14 | 1.00 |
| M3-2 | 49451.0 | 25.0 | 0.54 | 32.5 | 38.30 | 0.80 | 0.17 | 1.00 |
| M3-3 | 48017.0 | 21.0 | 0.40 | 21.5 | 22.28 | 0.87 | 0.13 | 1.00 |
| M4-1 | 48242.0 | 26.0 | 1.69 | 26.17 | 27.0 | 0.23 | 0.52 | 1.00 |
| M4-2 | 45542.0 | 28.0 | 1.75 | 31.0 | 33.64 | 0.22 | 0.53 | 1.00 |
| M4-3 | 51758.0 | 24.0 | 1.73 | 25.0 | 27.30 | 0.23 | 0.54 | 1.00 |
| M5-1 | 64877.0 | 24.0 | 0.24 | 25.0 | 27.22 | 0.92 | 0.08 | 1.00 |
| M5-2 | 48639.0 | 24.0 | 0.57 | 24.5 | 25.56 | 0.80 | 0.18 | 1.00 |
| M5-3 | 46373.0 | 24.0 | 0.49 | 24.5 | 25.36 | 0.84 | 0.16 | 1.00 |

**Table S2. Fungal diversity index analysis table for mask samples**

| Sample | Number | OTUs | Shannon | Chao | Ace | Simpson | Shannoneven | Coverage |
|---|---|---|---|---|---|---|---|---|
| M1-2 | 95200.0 | 81.0 | 1.41 | 91.11 | 92.24 | 0.42 | 0.32 | 1.00 |
| M1-3 | 60672.0 | 61.0 | 1.25 | 74.33 | 78.02 | 0.42 | 0.30 | 1.00 |
| M2-1 | 61923.0 | 61.0 | 1.80 | 70.33 | 66.62 | 0.27 | 0.44 | 1.00 |
| M2-3 | 63040.0 | 67.0 | 1.44 | 85.2 | 83.28 | 0.34 | 0.34 | 1.00 |
| M4-1 | 60638.0 | 66.0 | 1.33 | 79.33 | 81.66 | 0.47 | 0.32 | 1.00 |
| M4-2 | 76828.0 | 74.0 | 2.39 | 89.0 | 90.03 | 0.13 | 0.55 | 1.00 |
| M4-3 | 68677.0 | 82.0 | 1.76 | 89.33 | 88.36 | 0.25 | 0.40 | 1.00 |
| M5-1 | 63374.0 | 78.0 | 1.75 | 93.55 | 112.70 | 0.27 | 0.40 | 1.00 |
| M5-2 | 62763.0 | 22.0 | 1.35 | 22.0 | 22.74 | 0.34 | 0.44 | 1.00 |
| M5-3 | 61051.0 | 56.0 | 1.54 | 67.14 | 87.24 | 0.34 | 0.38 | 1.00 |

**Table S3. Bacterial diversity index analysis table for stage and human skin samples**

| Sample | Number | OTUs | Shannon | Chao | Ace | Simpson | Shannoneven | Coverage |
|---|---|---|---|---|---|---|---|---|
| Q1 | 80874.0 | 93.0 | 1.57 | 97.11 | 100.71 | 0.30 | 0.35 | 1.00 |
| Q2 | 85255.0 | 81.0 | 1.89 | 92.38 | 93.55 | 0.25 | 0.43 | 1.00 |
| Q3 | 72575.0 | 83.0 | 1.18 | 90.8 | 92.68 | 0.54 | 0.27 | 1.00 |
| C1 | 71121.0 | 115.0 | 1.70 | 120.2 | 122.74 | 0.40 | 0.36 | 1.00 |
| C2 | 69875.0 | 94.0 | 2.17 | 100.88 | 100.51 | 0.19 | 0.48 | 1.00 |
| C3 | 74450.0 | 122.0 | 1.71 | 130.5 | 133.69 | 0.38 | 0.36 | 1.00 |
| D1 | 67544.0 | 100.0 | 1.56 | 103.47 | 109.73 | 0.42 | 0.34 | 1.00 |
| D2 | 67716.0 | 137.0 | 2.22 | 143.18 | 144.68 | 0.17 | 0.45 | 1.00 |
| D3 | 67854.0 | 111.0 | 2.31 | 114.24 | 118.28 | 0.17 | 0.49 | 1.00 |
| F | 70008.0 | 102.0 | 2.12 | 114.0 | 113.90 | 0.23 | 0.46 | 1.00 |

**Table S4. Fungal diversity index analysis table for stage samples**

| Sample | Number | OTUs | Shannon | Chao | Ace | Simpson | Shannoneven | Coverage |
|---|---|---|---|---|---|---|---|---|
| Q1 | 51090.0 | 85.0 | 1.59 | 121.14 | 126.38 | 0.38 | 0.36 | 1.00 |
| Q2 | 62534.0 | 89.0 | 1.72 | 115.25 | 110.71 | 0.30 | 0.38 | 1.00 |
| Q3 | 52468.0 | 88.0 | 1.36 | 97.07 | 103.53 | 0.49 | 0.30 | 1.00 |
| C1 | 48949.0 | 105.0 | 2.21 | 124.25 | 140.40 | 0.19 | 0.48 | 1.00 |
| C2 | 63214.0 | 90.0 | 1.79 | 97.0 | 102.58 | 0.24 | 0.40 | 1.00 |
| C3 | 71357.0 | 83.0 | 2.48 | 84.5 | 84.72 | 0.15 | 0.56 | 1.00 |
| D1 | 88892.0 | 117.0 | 2.38 | 120.67 | 125.42 | 0.17 | 0.50 | 1.00 |
| D2 | 99165.0 | 136.0 | 2.38 | 149.0 | 145.13 | 0.22 | 0.48 | 1.00 |
| D3 | 101021.0 | 91.0 | 2.16 | 93.14 | 93.53 | 0.27 | 0.48 | 1.00 |

**Table S5. Molecular identification of bacterial strains isolated from mask, stage, and human skin samples**

| Nucleotide Blast reference strains | | | | |
|---|---|---|---|---|
| **Bacteria** | **Clost relative strain** | **Accession number** | **Phylum** | **Similarity(%)** |
| B1 | Exiguobacterium acetylicum strain | KT767813.1 | Firmicutes | 99.93% |
| B2 | Acinetobacter ursingii strain | MH113156.1 | Acinetobacter | 99.79% |
| B3 | Pseudomonas tructae strain | CP035952.1 | Proteobacteria | 100.00% |
| B4 | Stenotrophomonas rhizophila strain | KU921558.1 | Proteobacteria | 99.79% |
| B5 | Myroides odoratimimus strain | JF775419.1 | Bacteroidota | 100.00% |
| B6 | Alcaligenes faecalis strain | KX097967.1 | proteobacteria | 99.93% |
| B7 | Acinetobacter lwoffii strain | MN704528.1 | Acinetobacter | 99.58% |
| B8 | Bacillus cereus strain | MT605291.1 | Firmicutes | 99.52% |
| B9 | Pseudomonas fluorescens strain | OP341878.1 | Proteobacteria | 99.86% |
| B10 | Stenotrophomonas maltophilia strain | MT078668.1 | Proteobacteria | 100.00% |
| B11 | Pseudomonas rhizoryae strain | MK759856.1 | Proteobacteria | 99.43% |
| B12 | Pigmentiphaga litoralis strain | NR_044530.1 | Proteobacteria | 99.78% |
| B13 | Providencia manganoxydans strain | MH644827.2 | enterobacteria | 99.93% |

[a]The strains "B1 to B13" were identified by the 16S rDNA gene; all of them could be annotated to species.

**Table S6. Molecular identification of fungal strains isolated from mask and stage samples**

| Fungi | Clost relative strain | Accession number | Phylum | Similarity(%) |
|---|---|---|---|---|
| | Nucleotide Blast reference strains | | | |
| F1 | Meyerozyma guilliermondii strain | MG601180.1 | Ascomycota | 100.00% |
| F2 | Fusarium oxysporum strain | MT254999.1 | Ascomycota | 100.00% |
| F3 | Rigidoporus vinctus strain | MW077092.1 | Basidiomycota | 100.00% |
| F4 | Hannaella luteola strain | KP132260.1 | Basidiomycota | 100.00% |
| F5 | Aureobasidium pullulans strain | MH129527.1 | Ascomycota | 100.00% |
| F6 | Moesziomyces aphidis strain | MK994026.1 | Basidiomycota | 100.00% |
| F7 | Aureobasidium leucospermi strain | JN712487.1 | Ascomycota | 99.65% |
| F8 | Trichoderma atroviride strain | MH153634.1 | Ascomycota | 100.00% |
| F9 | Rhodotorula mucilaginosa strain | MT378424.1 | Basidiomycota | 99.83% |
| F10 | Exophiala xenobiotica strain | MZ573442.1 | Ascomycota | 99.17% |
| F11 | Penicillium citreonigrum strain | EU497944.1 | Ascomycota | 100.00% |
| F12 | Cladosporium parahalotolerans strain | MK796044.1 | Ascomycota | 100.00% |
| F13 | Penicillium citrinum strain | OP237250.1 | Ascomycota | 100.00% |
| F14 | Talaromyces verruculosus strain | OP237348.1 | Ascomycota | 99.82% |
| F15 | Sarocladium strictum strain | MK361137.1 | Ascomycota | 100.00% |
| F16 | Fusarium fujikuroi strain | MT549849.1 | Ascomycota | 100.00% |
| F17 | Cladosporium tenuissimum strain | OQ629133.1 | Ascomycota | 99.45% |

[a]The strains "F1 to F17" were identified by the ITS rDNA gene; all of them could be annotated to species.

**Table S7. The proportion of shared bacteria/fungi among the masks, stage, and the surface of insect eggs relative to the bacterial/fungal species on the surface of the insect eggs.**

| Bacteria | Proportion | Fungi | Proportion |
|---|---|---|---|
| Pseudomonas | 0.014 | Cladosporium | 0.4313 |
| Stenotrophomonas | 0.0046 | Alternaria | 0.1941 |
| Massilia | 0.0233 | Aspergillus | 0.22 |
| Sphingomonas | 0.532 | Fusarium | 0.6507 |
| Allorhizobium-Neorhizobium-Pararhizobium-Rhizobium | 0.0093 | g__norank_Eukaryota | 3.3035 |
| | | Kodamaea | 0.0148 |

**Figure S1. Determination of microbial community diversity in maske and stage samples using amplicon sequencing**

(A) Analysis of species diversity among mask samples at the OTU level based on 16S amplicon sequencing.(B) Analysis of species diversity among stage samples at the OTU level based on 16S amplicon sequencing.(C) Analysis of species diversity among mask samples at the OTU level based on 18S amplicon sequencing.(D) Analysis of species diversity among stage samples at the OTU level based on 18S amplicon sequencing.

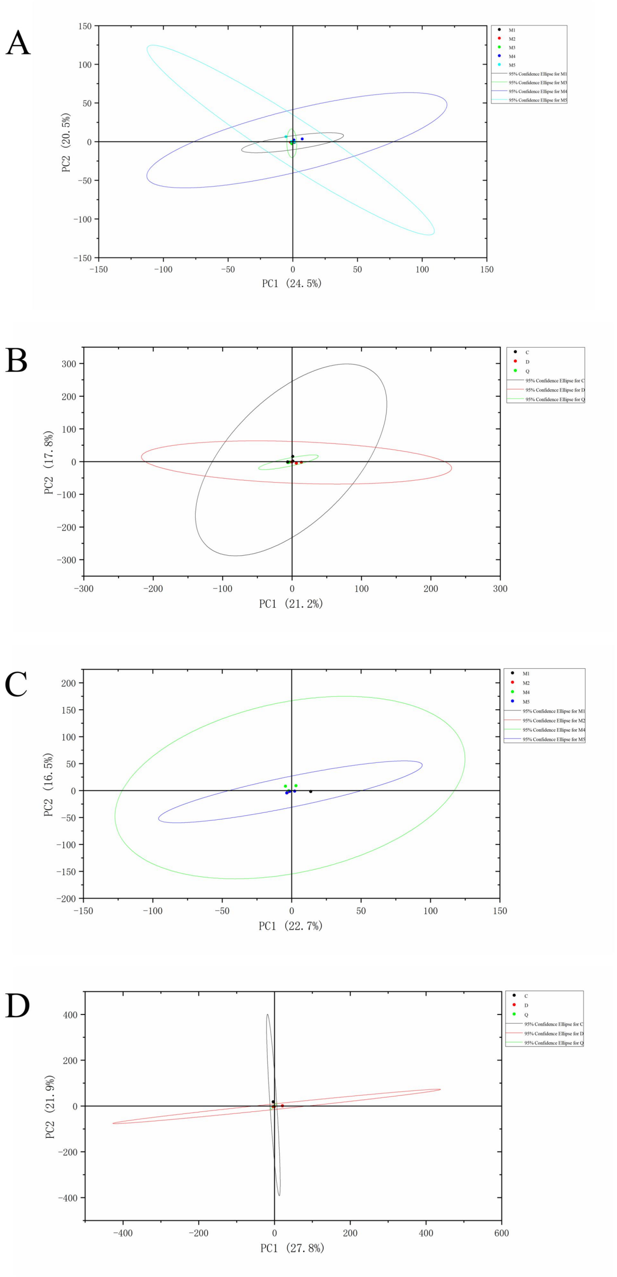


**Figure S2. Interaction Diagram of Microbial Communities in Wooden Heritage from Anshun Tunbao.**

In the interaction among microbial species on masks, stages, and the surrounding environment, firstly, the environmental microbes such as Pseudomonas, Stenotrophomonas, Aspergillus, Cladosporium, and Alternaria are transmitted to the masks and the stage through the movement of insects. The dominant microbe Aspergillus on the masks is then spread to the stage via insect activity. Similarly, the dominant microbe Stenotrophomonas on the stage is transmitted back to the masks through insects as well. This reciprocal transmission ultimately increases the diversity of microbes on the surfaces of the masks and the stage, expands the colonization range of these microbes, and exacerbates the deterioration of the wooden heritage.

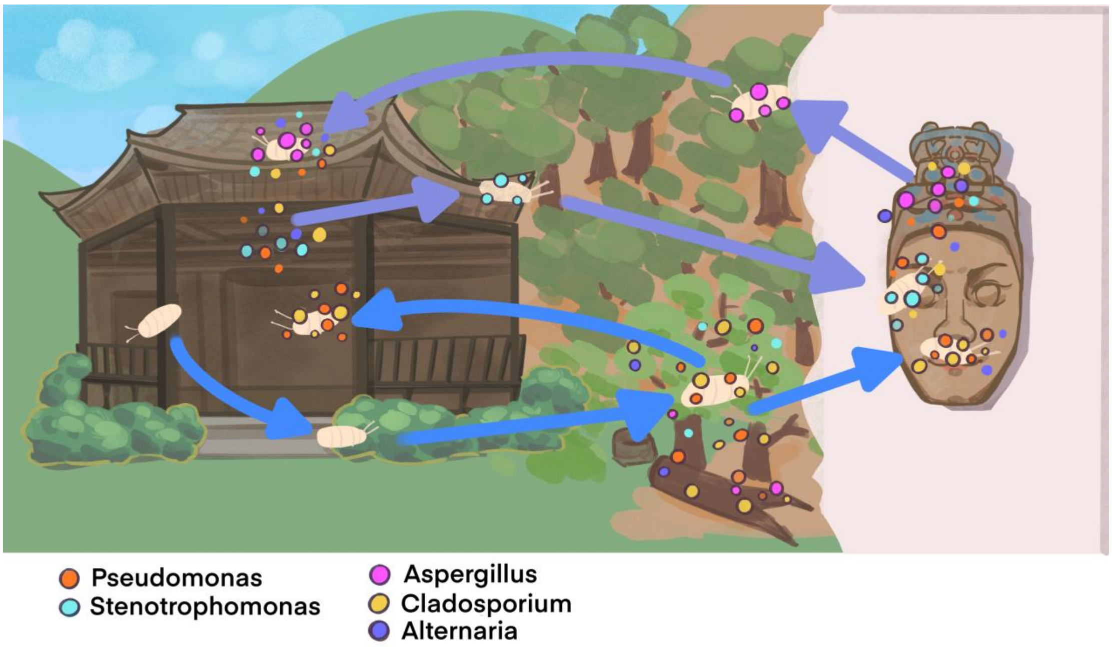
